# Shared book reading promotes experience-dependent autonomic synchrony in parent–preterm infant dyads

**DOI:** 10.64898/2026.05.19.726001

**Authors:** Laura Lavezzo, Ben Meuleman, Didier Grandjean, Edouard Gentaz, Sylvain Delplanque, Leonardo Ceravolo, Enzo Pasquale Scilingo, Petra Hüppi, Francisca Barcos-Munoz, Cristina Borradori-Tolsa, Mimma Nardelli, Manuela Filippa

## Abstract

Preterm birth is associated with alterations in early caregiver–infant regulation, with potential consequences for socio-emotional and physiological development. However, the mechanisms through which early interactional experience shapes these processes remain unclear. Here, we tested whether a structured dyadic intervention could modify co-regulatory dynamics across physiological, behavioral, and relational levels. Fifty-four 7-month-old preterm infants and their parents were assigned to either a shared book reading intervention (n = 22) or an active control condition based on a shared building activity (n = 32) and compared with 39 full-term infants. The intervention consisted of an 8-week program of shared book reading, designed to structure parent–infant interaction. Physiological synchrony was assessed at the dyadic level, alongside infants’ autonomic regulation and cardiovascular signal complexity. Behavioral engagement and parental attachment representations were also evaluated.

Results showed that mother–infant physiological synchrony emerged selectively within the interactional context trained by the intervention and only in the intervention group. This context-specific synchrony was accompanied by modulation of vagal activity and increased cardiovascular complexity in preterm infants, consistent with enhanced flexibility of autonomic control. At the behavioral and relational levels, intervention infants showed increased initiating joint attention, while parents reported higher secure attachment.

These findings support a model of experience-dependent early synchrony, in which repeated dyadic interaction through shared book reading shapes the coupling between interpersonal coordination and individual physiological regulation. By linking synchrony, autonomic flexibility, and social engagement, this study identifies a mechanism through which early caregiving experience can organize developmental trajectories following prematurity.

## Introduction

Early caregiver–infant interactions play a central role in shaping physiological regulation and socio-emotional development across the first years of life. Converging evidence shows that sensitive caregiving supports the maturation of stress-regulatory systems, including cortisol rhythms and vagal control, while adverse or inconsistent interactions can disrupt these processes and increase vulnerability (1–4). At the neural level, early relational experiences become biologically embedded within corticolimbic circuits involved in emotion regulation during sensitive developmental windows (5), while early dyadic exchanges contribute to the development of infant’s socio-emotional competences (6–8).

These processes may be particularly vulnerable following preterm birth, which is associated with altered trajectories of autonomic and relational functioning. Preterm infants often exhibit reduced parasympathetic activity and atypical heart rate variability early in life, reflecting disruptions in autonomic maturation (9–11) and lower levels of autonomic synchrony (12). Such alterations can persist across development and are accompanied by increased risk for socio-emotional difficulties and differences in caregiver–infant interaction patterns (13–15). Importantly, early interventions targeting parent–infant interaction have been shown to support both autonomic regulation and relational functioning in preterm populations (16–19).

These findings underscore the importance of identifying mechanisms that support early interaction between parents and preterm infants. Within this framework, co-regulation between caregiver and infant is thought to rely on dynamic coordination across behavioral and physiological systems. In particular, physiological synchrony, defined as the temporal alignment of biological signals between individuals, has been proposed as a key mechanism supporting early regulation (20).

However, the extent to which physiological synchrony reflects a dynamic process shaped by early relational experience remains an area for further research. Existing evidence suggests that synchrony is highly context-sensitive, varying across interactional demands, affective states, and environmental conditions, which points to a strong state-dependent component rather than a fixed dyadic trait (21–23). Consistently, emerging evidence suggests that synchrony can be shaped by early interventions. Interactional contexts involving touch, proximity, and contingent engagement have been shown to modulate neural and, in some cases, autonomic synchrony, and these effects are linked to caregiving quality and developmental outcomes (24–27). Longitudinal findings further indicate that early caregiving experiences can have enduring effects on dyadic coordination, consistent with experience-dependent tuning of interpersonal processes (26, 28). Early co-regulation is thus experience-dependent, emerging through repeated dyadic exchanges that are structured by specific relational contexts. Longitudinal evidence indicates that forms of co-regulation evolve across infancy through repeated interaction, and are associated with later attachment security and developmental outcomes (29–31). Importantly, co-regulation varies across interactional settings, with different contexts selectively shaping the organization of dyadic exchanges (20, 32, 33).

Within this framework, synchrony should not be considered a simple byproduct of shared activity, but rather a property that is selectively shaped by the temporal and structural features of interaction. We therefore hypothesize that specific interactional formats may differentially support the emergence of coordinated physiological and behavioral processes.

Shared book reading represents one such context, providing a structured yet flexible interactional environment in which adult and infant jointly attend to a common object, and where the organization of turns, attention, and affect is scaffolded by the book itself (34–36). By providing a shared focus and predictable interactional framework, book reading can facilitate reciprocal exchanges and sustained engagement (37, 38) while allowing caregivers to flexibly adapt their behavior to the infant’s signals (39). Importantly, shared reading has been associated with increased parental sensitivity, reduced stress, and enhanced infant engagement, including joint attention and communicative participation (37, 40, 41). A substantial body of empirical and meta-analytic research demonstrates that shared book reading is consistently associated with gains in expressive and receptive language, with meta-analyses reporting small-to-moderate effects on vocabulary and broader language outcomes (42, 43). Longitudinal studies further show that early exposure to book reading is associated with later vocabulary, grammar, and reading comprehension, even after controlling for socioeconomic status and general parental language input (44–46). Beyond language, shared reading has been linked to broader cognitive and socio-emotional development, including social competence, attention, and reduced behavioral difficulties (47–49). These effects appear to be particularly driven by interactive and dialogic reading practices, which promote joint attention, communicative signaling, and emotional understanding (50–53).

However, to our knowledge, randomized clinical trials examining the effects of shared book reading interventions in early infancy are scarce (37), and studies investigating their underlying mechanisms are currently lacking, particularly in at-risk populations such as preterm infants.

Building on this gap, we examined whether a shared book reading intervention could influence parent–infant autonomic synchrony and individual autonomic, behavioral, and relational outcomes. Late preterm infants at 7 months of corrected age and their caregivers were assigned to either a shared book reading intervention or an active control condition involving shared play and compared with full-term dyads. By integrating measures of dyadic physiological coupling, infants’ autonomic regulation and cardiovascular complexity, early social-communicative behavior, and parental attachment representations, the present study aimed to investigate whether early shared book reading intervention can shape co-regulation across multiple levels of functioning.

## Results

### Overview of infant demographic data

#### Population

A total of 54 seven-month-old preterm infants (gestational age: 33.80 ± 0.98 weeks, range 31.57–36.14; corrected age at testing: 38.22 ± 3.17 weeks, range 31.71–45.57) and their parents participated in the study. Of these, 22 infants were assigned to the Shared Book Reading activity (PI, Preterms Intervention), while 32 infants participated in a Building block Shared activity (PC, Preterms active Control). In addition, a passive control group consisting of 39 nine-month-old full-term infants (T, Terms) was recruited for comparison.

#### Preterm infants’ demographic comparison

Characteristics of preterm infants in the two groups have been compared, results are reported in Table 2. No significant differences were observed between the intervention and active control groups for any baseline characteristic (all p > 0.05).

**Table 1.** Demographic characteristics of infants by group. Values are reported separately for each group.

| Characteristic | PI, N = 22<sup>1</sup> | PC, N = 32<sup>1</sup> | T, N = 39<sup>1</sup> |
| --- | --- | --- | --- |
| **Sex** |  |  |  |
| M | 13 | 17 | 26 |
| F | 9 | 15 | 13 |
| **Age at T0 (weeks)** | $35.57 \pm 1.80$ | $35.26 \pm 1.37$ | / |
| **Age at T1 (weeks)** | $45.09 \pm 3.46$ | $45.81 \pm 2.84$ | $43.30 \pm 3.50$ |
| **Age at T0 Corrected (weeks)** | $28.39 \pm 2.04$ | $28.18 \pm 1.49$ | / |
| **Age at T1 Corrected (weeks)** | $38.07 \pm 3.50$ | $38.34 \pm 2.98$ | $43.30 \pm 3.50$ |
| **Apgar 1 min** | $6.71 \pm 2.95$ | $7.61 \pm 2.36$ | $6.56 \pm 3.12$ |
| **Apgar 5 min** | $8.43 \pm 1.57$ | $8.97 \pm 1.11$ | $8.15 \pm 2.30$ |
| **Apgar 10 min** | $9.10 \pm 1.18$ | $9.45 \pm 0.89$ | $9.09 \pm 1.36$ |
| **Gestational Age (weeks)** | $33.82 \pm 1.05$ | $33.79 \pm 0.95$ | $39.72 \pm 1.35$ |
| **Weight at birth (gr)** | $2,172.95 \pm 372.32$ | $2,005.00 \pm 412.33$ | $3,243.97 \pm 457.04$ |
| **Cranial perimeter (cm)** | $31.11 \pm 1.63$ | $30.52 \pm 1.64$ | $34.99 \pm 3.39$ |
| **Percentile w** | $2.23 \pm 0.75$ | $2.00 \pm 1.05$ | $2.09 \pm 0.98$ |
| **Percentile p** | $2.18 \pm 0.85$ | $1.78 \pm 1.01$ | $1.94 \pm 0.93$ |
| **SES** | $4.63 \pm 2.55$ | $4.60 \pm 2.37$ | $4.00 \pm 2.34$ |
| **Mother's age (year)** | $33.50 \pm 6.52$ | $34.63 \pm 4.95$ | $33.00 \pm 5.18$ |

<sup>1</sup>n; Mean ± SD

**Table 2.**
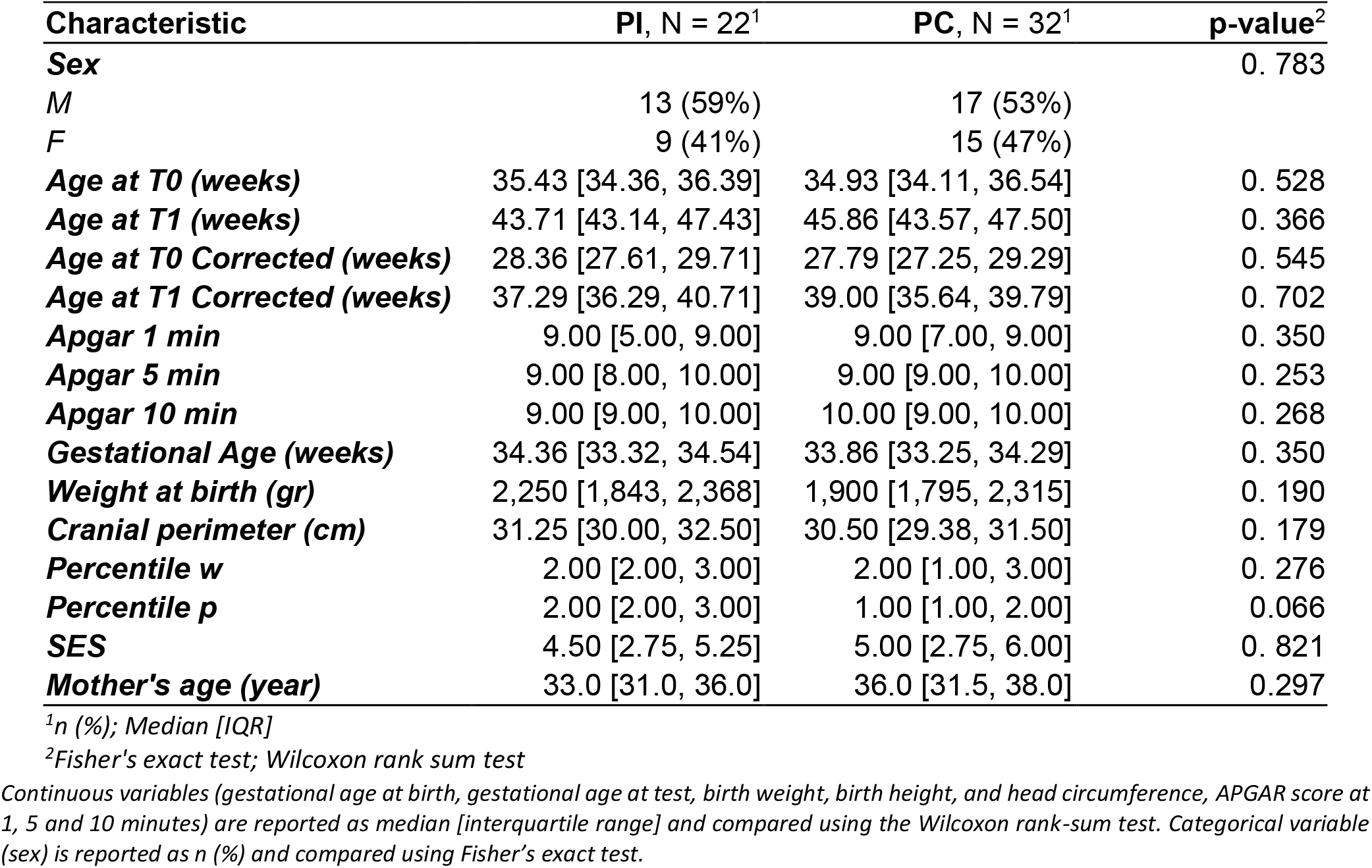
Demographic comparison of preterm infants.

#### Preterm infants’ parental report comparison at T0

*F*-tests were conducted to examine baseline differences between the preterm intervention and control groups in outcome measures, using linear regression models that adjusted for propensity scores. No significant group differences were observed for any outcome measure, including PSI Total Stress (*F*(1,19) = 0.343, *p* = .57, η^2^ =0,015), PSI subscales (all *p* ≥ .31, all η^2^ < 0.40), RSQ Total and subscales (all *p* ≥ .29, all η^2^ < 0.54), or EPDS (*F*(1,19) = 0.036, *p* = .85, η^2^ <0.01). These results indicate that the two groups were comparable on all parent’s psychological measures prior to the intervention.

### SHER affects autonomic synchrony

To test mother-infant autonomic synchrony, a total of 62 mother-infant dyads had usable ECG data for the interactional tasks (*Preterm Intervention* = 15; *Preterm Control* = 18; *Term* = 29). Analysis has been done on the total task window (3-min play, 3-min reading).

Mother–infant physiological coupling was assessed separately for respiratory and residual components, after *z*-score normalization. This decomposition was essential because respiratory variability during the reading task could influence frequency-based autonomic and synchrony metrics. Synchrony was quantified using Cross-Recurrence Quantification Analysis (CRQA), with the Rate of Determinism (DET) as the primary metric and tested against surrogate dyads (i.e. shuffled mother-infant pairs) to determine whether observed coupling exceeded chance. Because significant synchrony emerged only in specific groups and task phases, no ANOVA model was applied.

For the respiratory component, significant determinism, reflecting a structured respiratory coupling between mother and infant, was observed in the Intervention group during the reading phase (KS test, *D* = 0.446, *p* = 0.002). No other group or phase comparisons reached significance (all *p* > 0.25, *D* < 0.19).

For the residual component, significant determinism was detected in the Term group during both play (*D* = 0.254, *p* = 0.020) and reading (*D* = 0.305, *p* = 0.005), whereas no other groups or phases showed significant coupling (all *p* > 0.66, *D* < 0.11).

**Figure 1.**
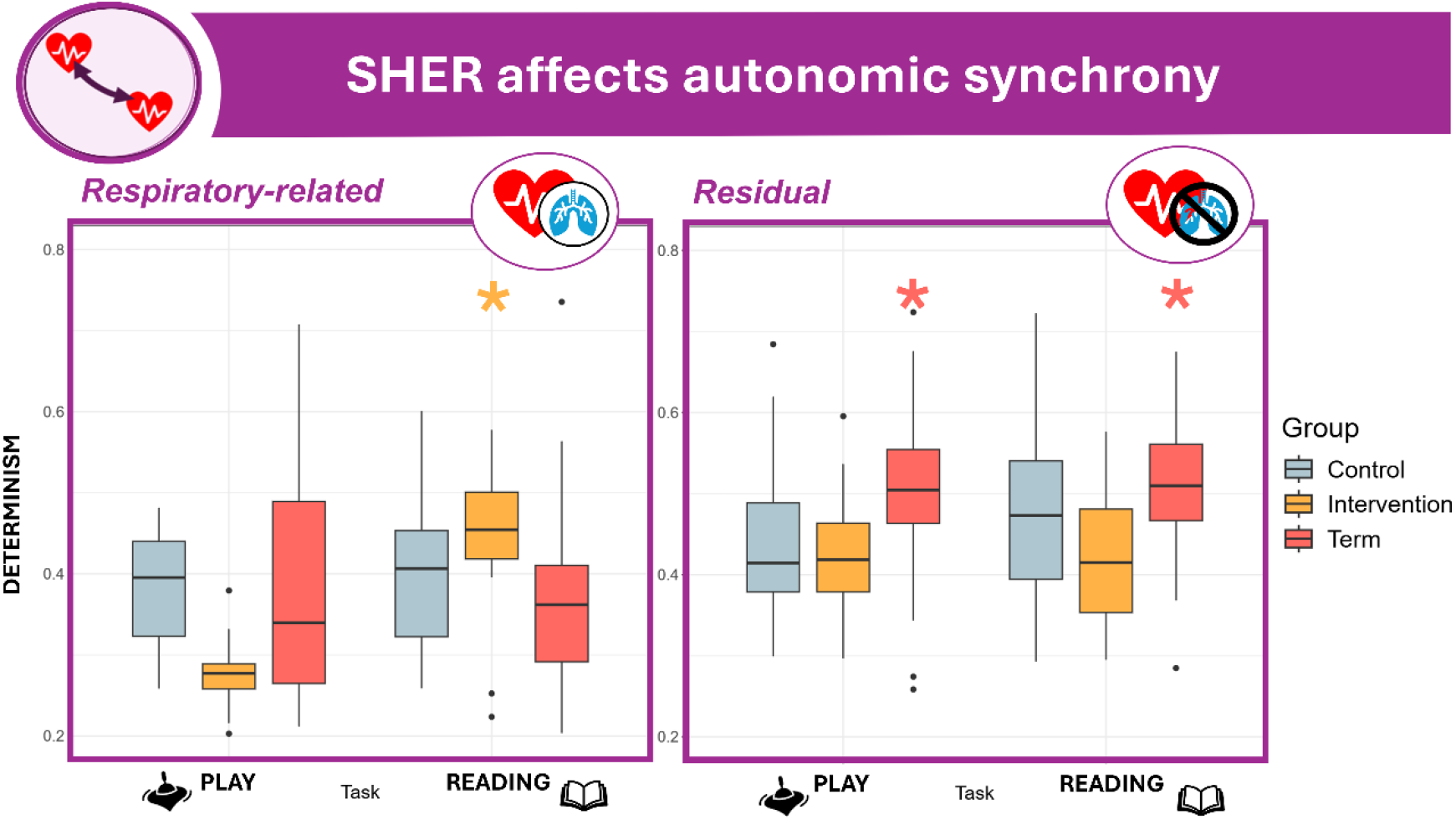
Autonomic synchrony indices across experimental tasks. Boxplots show the median and interquartile range for autonomic synchrony (Determinism rate from CRQA) for both the respiratory-related HRV component (left plot) and the residual HRV component (right plot). The x-axis indicates task (Play, Reading). Infant groups are colour-coded (light blue, preterm control; yellow, preterm intervention; red, term passive control). Asterisks, color-coded by group, denote significant synchrony (P < 0.05).

### SHER modulates autonomic reactivity

Having characterized mother–infant physiological synchrony at the dyadic level, we next examined whether these interactional patterns were accompanied by changes in infants’ individual autonomic regulation across task contexts.

A total of 70 infants had usable ECG data (Preterm Intervention = 16; Preterm Control = 21; Term = 33). Parent– infant dyads completed four consecutive sessions (1-min baseline, 3-min play, 3-min reading, 1-min final baseline) in a secluded setting with minimal experimenter interference. A 1-min analysis window was used across conditions to match baseline durations following established Heart Rate Variability (HRV) standards (54).

#### Vagal activation

Parasympathetic activity was quantified using time-domain (RMSSD), frequency-domain (HF power, 0.2–1.5 Hz), and nonlinear (SD1) HRV indices.

Mixed effects ANOVAs showed no significant main effect of Group for any HRV index (*all p* > 0.6), indicating comparable overall vagal tone across groups. In contrast, a significant main effect of Task was observed for RMSSD (*F*(3, 193.16) = 3.49, *p* = 0.017, *η*^*2*^ = 0.06), HF power (*F*(3, 193.74) = 3.14, *p* = 0.026, *η*^*2*^ = 0.05), and SD1 (*F*(3, 193.16) = 3.49, *p* = 0.017, *η*^*2*^ = 0.06), reflecting within-session modulation of vagal activity. No significant Group × Task interaction was detected (*all p* > 0.23).

#### Post-hoc contrasts

While the overall ANOVAs did not reveal significant group differences, exploratory post-hoc analyses were examined descriptively to identify potential trends of clinical interest, that may not emerge in multifactorial models with relatively small samples.

Within-group comparisons revealed that in the Intervention group, HRV indices significantly decreased from the initial baseline to the play phase: RMSSD (*t*(193) = 2.95, *p* = 0.019, *d* = 1.04), HF power (*t*(193) = 2.67, *p* = 0.041, *d* = 0.94) and (SD1: *t*(193) = 2.95, *p* = 0.019, *d* = 1.04).

Additionally, the HF power was significantly lower in the reading phase than the baseline for the Intervention group only (*t*(193) = 2.89, *p* = 0.022, *d* = 1,02).

No significant within-task differences were observed for the Control or Term groups (*all p* > 0.3).

While task context influenced HRV overall, significant autonomic modulation emerged exclusively in the SHER intervention group, which exhibited decreased vagal activity during shared play. No significant changes were detected in either control group.

#### Complexity assessment

Cardiovascular signal complexity reflects the adaptability and flexibility of autonomic control, with higher values indicating a richer, more variable temporal structure consistent with a system capable of dynamically responding to internal and external demands.

Cardiovascular signal complexity was examined using Sample Entropy (SampEn) and the Complexity Index (CI), computed as the area under the Fuzzy Entropy curve across the first three coarse-grained scales (Costa et al., 2002).

Mixed effect ANOVAs showed a significant main effect of Group for both SampEn (*F*(2, 65.01) = 4.83, *p* = 0.011, *η*^*2*^ = 0.13) and CI (*F*(2, 64.79) = 3.27, *p* = 0.045, *η*^*2*^ = 0.09), indicating overall differences in cardiovascular complexity among groups. SampEn also varied across Task phases (*F*(3, 194.5) = 3.43, *p* = 0.018, *η*^*2*^ = 0.07), suggesting phase-dependent modulation. The Group × Task interaction approached significance for SampEn (*F*(6, 194.71) = 2.04, *p* = 0.062, *η*^*2*^ = 0.06) and was significant for CI (*F*(6, 194.50) = 2.17, *p* = 0.046, *η*^*2*^ = 0.06), warranting post-hoc examination of group-specific changes.

#### Post-hoc contrasts

Exploratory post-hoc comparisons revealed within-session modulation of complexity: in the intervention group SampEn significantly increased from the initial baseline to the reading phase (*t*(193) = −3.65, *p* = 0.002, *d* = −1.29), and CI showed a similar increase (*t*(193) = −3.31, *p* = 0.006, *d* = −1.16). In the Control group, SampEn significantly increased from the initial baseline to the play phase (*t*(193) = −2.80, *p* = 0.028, *d* = −0.86). No other within-group contrasts were significant.

Between-group post-hoc comparisons indicated that both metrics were significantly higher in the Intervention group compared with both Control and Term infants. Specifically, for SampEn, values were greater in the Intervention group than Control (*t*(194) = −2.54, *p* = 0.032, *d* = −1.02) and Term (*t*(161) = 3.86, *p* = 0.0005, *d* = 1.58) during the reading. Whereas during the Play, Term had lower values than both preterm Control (*t*(155) = 2.72, *p* = 0.020, *d* = 1.04) and Intervention (*t*(161) = 2.66, *p* = 0.031, *d* = 1.04).

Similarly, for CI, the Intervention group showed higher values than Control (*t*(195) = −2.59, *p* = 0.028, *d* = −1.04) and Term (*t*(162) = 3.73, *p* = 0.0008, *d* = 1.52).

**Figure 2.**
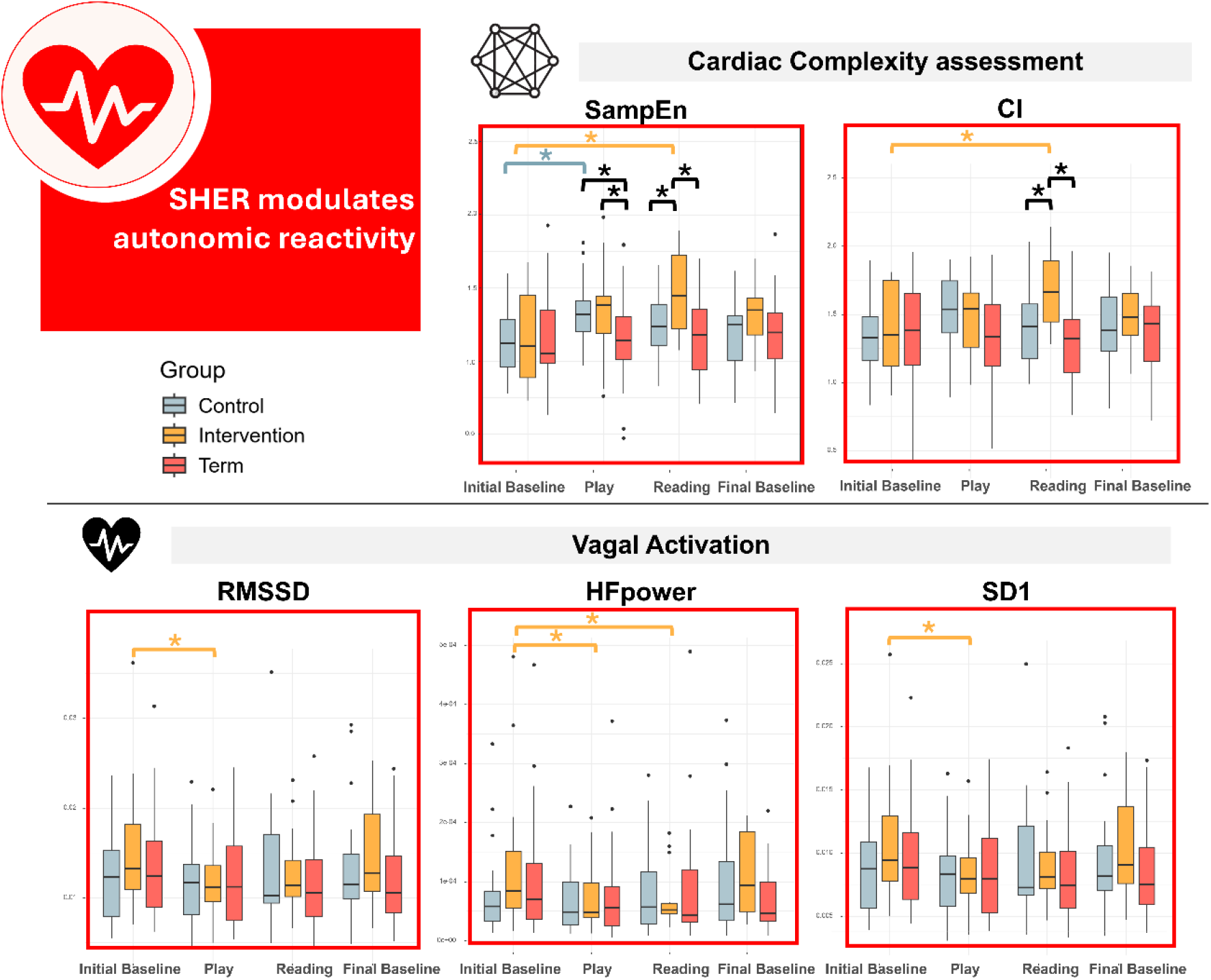
Autonomic complexity indices across experimental tasks (Upper plots). Boxplots show the median and interquartile range for Sample Entropy (SampEn) and Complexity Index (CI). The x-axis indicates task (Initial Baseline, Play, Reading, Final Baseline). **Parasympathetic indices across experimental tasks** (Lower Plots). Boxplots show the median and interquartile range for RMSSD, HF power, and SD1. The x-axis indicates task (Initial Baseline, Play, Reading, Final Baseline).Infant groups are colour-coded (light blue, preterm control; yellow, preterm intervention; red, term passive control). Asterisks denote significant differences (P < 0.05). Inter-task comparisons are colour-coded by group, whereas inter-group comparisons are shown in black.

#### SHER increase Infant- Initiated Joint Attention

A total of 50 infant behaviours has been scored after the intervention (Intervention = 15; Preterm Control = 15; Term = 20) using the Early Social-Communication Scales (ESCS). A significant main effect of Group (*F*(2, 45) = 3.50, *p* = 0.039, *η*^*2*^ = 0.13) emerged for Joint Attention Behaviours (IJA) refer to the child’s skill in employing nonverbal behaviours to involve others in the experience of objects or events. Post-hoc assessment revealed that Frequency with which a child uses eye contact, pointing and showing to initiate shared attention to objects or events was significantly more frequent in the Intervention group compared with Preterm Control infants (*t*(45) = 2.52, *p* = 0.04, *d* = 0.9). No significant effects were observed for the remaining scales (all p > .05).

#### Parents reported higher levels of secure attachment

Parent-infant relationship and attachment style had been investigated using the Relationship Scale Questionnaire is a self-reported questionnaire made by Griffin & Bartholomew in 1994. A total of 76 dyads data has been collected (Intervention = 19; Preterm Control = 22; Term = 35)

A significant main effect of Group (*F*(2, 71) = 6.69, *p* = 0.0022, *η*^*2*^ = 0.10) emerged for Secure attachment style. Post-hoc assessment revealed that Parents of preterm infant in the intervention group rated the Secure attachment with significantly higher values than Control (*t*(71) = 2.63, *p* = 0.028, *d* = 0.83) and Term (*t*(71) = 3.53, *p* = 0.002, *d* = 1.26). No significant effects were observed for the remaining scales (all *p* > .05).

**Figure 3.**
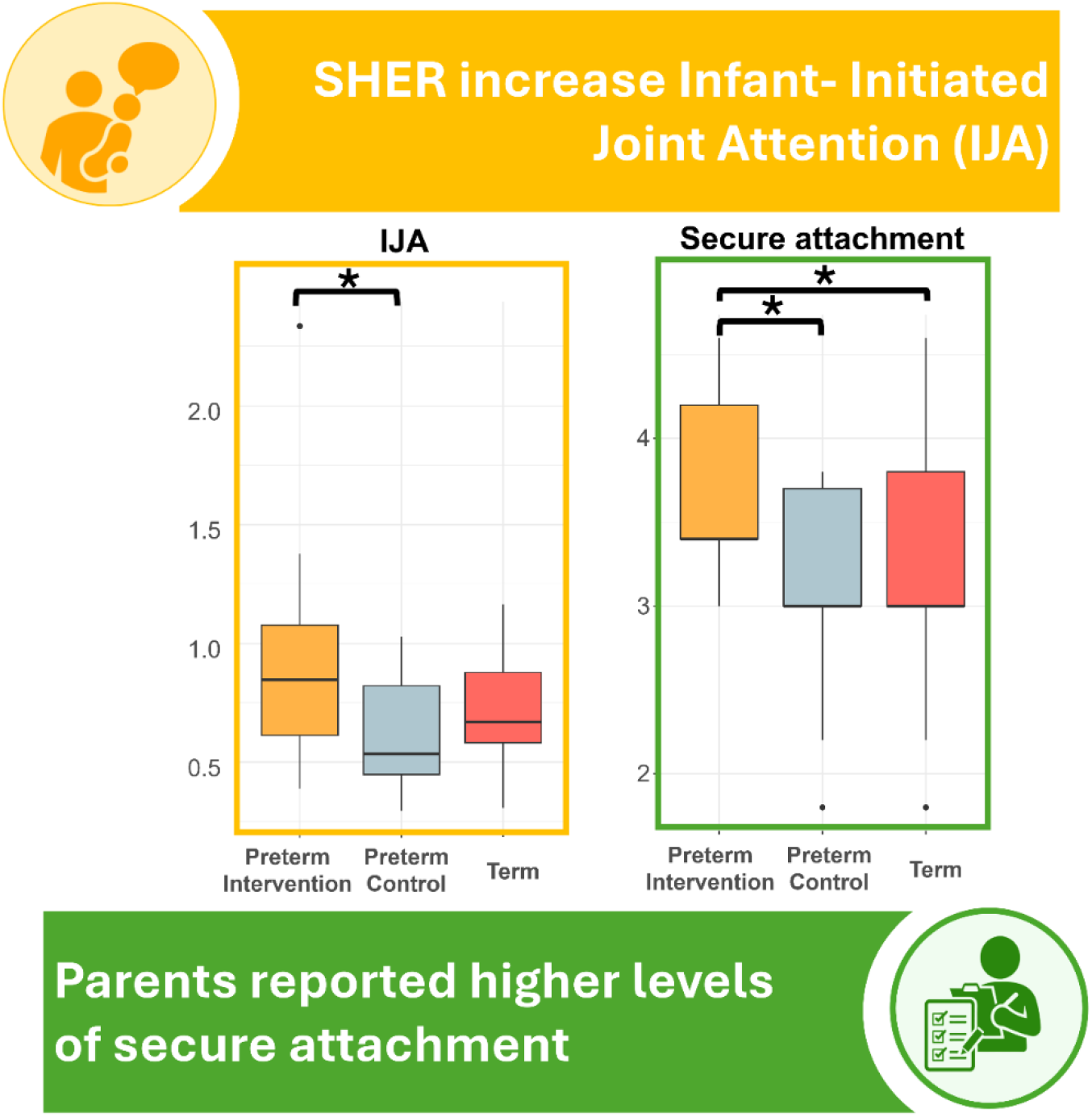
Baseline Infants’ Initiated Joint Attention behaviours (left plot) and parental self-evaluation (right plot). Boxplots show the median and interquartile range. Infant groups are colour-coded (light blue, preterm control; yellow, preterm intervention; red, term passive control). Asterisks denote significant inter-group differences (P < 0.05).

## Discussion

The present findings converge across dyadic, physiological, behavioral, and relational levels to suggest that the shared book reading intervention selectively shapes early co-regulatory processes in preterm infants. More broadly, these converging effects can be interpreted within an embodied framework of affective and cognitive development, which posits that physiology, behavior, and relationship quality are deeply intertwined and co-constructed through real-time caregiver–infant interaction (55–59). From this perspective, increased autonomic flexibility, enhanced parent–infant synchrony, greater infant-initiated joint attention, and higher parental reports of secure attachment are not independent outcomes, but coordinated expressions of a common developmental process. Dynamic systems account further suggest that changes at one level, such as physiological regulation, may enable new forms of engagement at others, thereby creating mutually reinforcing developmental cascades (59).

The present results confirm that at a dyadic level, parent–infant autonomic synchrony was specifically observed within the context structured by the intervention. This indicates that synchrony is not simply elicited by joint activity per se but depends on the temporal and relational organization of the interaction (20).

Shared book reading represents a particularly potent context in this respect, as it inherently organizes interaction through rhythmic vocalization, predictable turn-taking, and a shared attentional focus. Conversational and narrative structures impose temporal regularities, such as phrasing, pauses or repetitions that scaffold coordinated exchanges and aligning interpersonal timing, thereby potentially facilitating coordination across partners (60). In parallel, the joint focus on the book promotes sustained shared attention and engagement, which have been linked to increased neural and affective synchrony during narrative processing (61, 62). Crucially, this temporal and attentional organization is embedded within a bodily interactional frame. Shared reading with young infants typically involves close physical proximity, often with the infant seated on the caregiver’s lap, and creating conditions for postural alignment, shared breathing space, and tactile contact. Growing evidence indicates that such embodied co-presence plays a key role in shaping physiological regulation: affectionate touch and close contact can directly modulate infants’ autonomic activity, increasing parasympathetic tone and stabilizing cardiovascular rhythms (63, 64). In other dyadic contexts, physical touch has been shown to enhance autonomic and cardio-respiratory coupling, suggesting that bodily contact may provide a somatovisceral substrate for interpersonal synchrony (65–68). At the same time, proximity alone is not sufficient, as synchrony depends on the combination of physical closeness with active, contingent interaction (27). In this sense, shared book reading can be considered a prototypical embodied routine, in which gaze, posture, touch, vocalization, and affective exchange jointly organize interaction and provide repeated opportunities for co-regulation (69). These multimodal exchanges may scaffold the infant’s developing regulatory systems by linking bodily states with social signals and expectations, thereby supporting the gradual intersubjective capacities in at-risk populations (70).

Importantly, by dissociating respiratory and residual components, we observed a clear differentiation in how synchrony manifested across groups and contexts. Structured respiratory coupling was observed selectively in the intervention group during the reading phase, and synchrony in the residual component, less directly driven by respiratory dynamics, was observed in the term group across both play and reading, but not in preterm infants. This pattern suggests that residual coupling may index a more intrinsic form of autonomic co-regulation, reflecting the capacity of the dyad to achieve coordinated physiological regulation beyond immediate interactional constraints.

In this context, the residual component of heart rate variability (HRV) could reflect autonomic dynamics that are not directly driven by breathing and instead capture broader sympathetic and vagal influences (71–73). Such respiration-independent HRV signals could index central–autonomic integration within the central autonomic network, linking physiological regulation to affective and cognitive processes (74, 75). In this context, residual coupling between partners may reflect deeper, intrinsic regulatory organization in the term dyads (76, 77), often reported as affected by prematurity. The absence of this form of coupling in preterm dyads, even following intervention, may indicate that deeper autonomic co-regulatory processes are less readily shaped in the short term, or require more prolonged or developmentally mature interactions to emerge.

At the individual level, the emergence of context-specific synchrony was paralleled by marked modulation of infants’ autonomic function, including changes in vagal activity and increased cardiovascular complexity. These patterns are consistent with enhanced physiological flexibility, a core property of adaptive regulatory systems. Cardiovascular complexity reflects the integration of multiple control mechanisms operating across time scales, and higher complexity is generally interpreted as an index of a system capable of dynamically adjusting to changing environmental demands (78, 79). In early development, such flexibility is particularly critical, as infants rely on rapidly shifting physiological states to support attention, social engagement, and stress regulation.

This last aspect is especially relevant for preterm infants, who are known to exhibit altered autonomic maturation, including reduced heart rate variability and less stable cardiovascular control, reflecting a diminished capacity for adaptive regulation (80, 81). Consistently, these early physiological vulnerabilities extend to the behavioral level, as children born preterm frequently experience challenges in emotion and stress regulation, as well as reduced sustained attention, suggesting broader disruptions in emerging self-regulatory processes (82–84).

Within this framework, the increase in cardiovascular complexity observed in the preterm intervention group, together with higher autonomic synchrony with parents, suggests a shift toward a more flexible and responsive autonomic system, supporting greater engagement with the environment and social partners. These effects are consistent with previous work showing that shared reading interventions can enhance autonomic flexibility and physiological synchrony, particularly when caregivers are attuned to infant cues (55, 85). More generally, shared reading appears to provide a structured context in which improvements in physiological regulation may directly support orienting, sustained attention, and emotion regulation, thereby linking bodily flexibility to emerging cognitive and affective capacities (56, 86).

Importantly, this enhanced physiological flexibility and enhanced parent-infant synchrony were paralleled by converging behavioral and relational changes: infants in the shared reading group showed increased initiating joint attention, indicating greater engagement in shared social experiences, while parents reported higher levels of secure attachment. Rather than representing independent outcomes of the shared book reading intervention, these last effects likely reflect different expressions of a common underlying process, whereby improved autonomic regulation supports the infant’s capacity to orient, engage, and coordinate with the caregiver in real time. In this sense, higher levels of autonomic synchrony and increased cardiovascular complexity may provide the physiological substrate for the early emergence of social-communicative competence. This interpretation is supported by studies showing that infant-initiated joint attention is closely linked to regulatory capacity and embodied coordination during interaction (55, 87–89), and that relational gains following shared reading are accompanied by higher parental warmth, lower parenting stress, and stronger attachment-related processes (37, 57, 90). Intervention studies further suggest that improvements in caregiver–infant reciprocity during book sharing mediate gains in language, attention, and socio-emotional functioning, indicating that the quality of interaction acts not as a parallel correlate, but as a core mechanism linking shared routines to developmental change (70).

## Limitations

Several limitations should be acknowledged. Firstly, although the findings support a multi-level model of co-regulation, the underlying neural mechanisms remain untested. Future studies combining neuroimaging and hyperscanning approaches will be essential to characterize brain-to-brain and brain–heart coupling underlying these effects (20). Second, the extent to which these findings generalize across cultural contexts, caregiving practices, and different at-risk populations remains unclear, as co-regulatory processes are likely shaped by broader sociocultural environments. Finally, while shared book reading represents a structured and scalable intervention, questions remain regarding its specificity relative to other dyadic routines and the fidelity with which such interventions can be implemented in real-world settings. Addressing these gaps will be critical to clarify the mechanisms, generalizability, and long-term impact of early relational interventions.

## Conclusion

This study shows that early shared book reading can engage core mechanisms of co-regulation in preterm infants. Physiological synchrony emerged selectively within the interactional context shaped by the intervention and was accompanied by increased cardiovascular complexity, enhanced infant social engagement, and higher parental perceptions of secure attachment. Because these converging effects were observed only in the intervention group, they suggest that structured dyadic experience can simultaneously influence interpersonal coordination and individual autonomic regulation.

These findings support an experience-dependent view of early co-regulation, in which repeated caregiver–infant interaction shapes regulatory processes within specific relational contexts. From a clinical perspective, shared book reading may represent a feasible, low-intensity early intervention for preterm infants, with the potential to support physiological flexibility and parent–infant engagement during a sensitive developmental period.

## Significance Statement

Preterm birth can disrupt early caregiver–infant synchrony, altering developmental trajectories. This study introduces a model of experience-dependent early synchrony process, showing that targeted dyadic interaction can shape physiology, behavior, and relational experiences. A shared book reading intervention selectively promoted mother–infant physiological synchrony within the trained interactional context, alongside increased cardiovascular complexity, consistent with enhanced flexibility of autonomic control, infant social engagement and higher parental perceptions of secure attachment. These converging effects, observed only in the intervention group, indicate that early relational experience can shape interindividual synchrony in a context-dependent manner. By linking dyadic interaction with individual autonomic regulation, the findings identify a pathway through which early caregiving experience can organize development, highlighting the potential of targeted relational interventions to potentially modify outcomes following prematurity.

## Materials and Methods

### Population

#### Inclusion and exclusion criteria

For the preterm group, only infants born at 32 weeks of gestational age (GA) or earlier and without severe neurological lesions were included. For the term group, infants born at 37 weeks GA or later and without severe neurological lesions were eligible. Exclusion criteria included maternal drug or alcohol abuse during pregnancy. Parents of preterm infants were invited to attend an initial visit when their child reached 7 months of corrected age. Following informed consent, adequate time for consideration, and signing of consent forms, one parent was designated as the project’s responsible caregiver. Premature infants were then enrolled at 7 months corrected age and randomly assigned to either the intervention (Preterm Intervention, PI) or the control group (Preterm Control, PC). Term infants were exclusively included and evaluated at nine months (group T).

The experimental protocol was approved by the Geneva Cantonal Research Ethics Committee (CCER) in June 2021 (ID 2021-00528). The research was conducted in compliance with the ethical standards set out in the Declaration of Helsinki, the principles of good clinical practice, the Human Research Act (HRA), the Ordinance on Research Involving Human Subjects with the Exception of Clinical Trials (HRO), and other laws in force in Switzerland.

**Figure 4.**
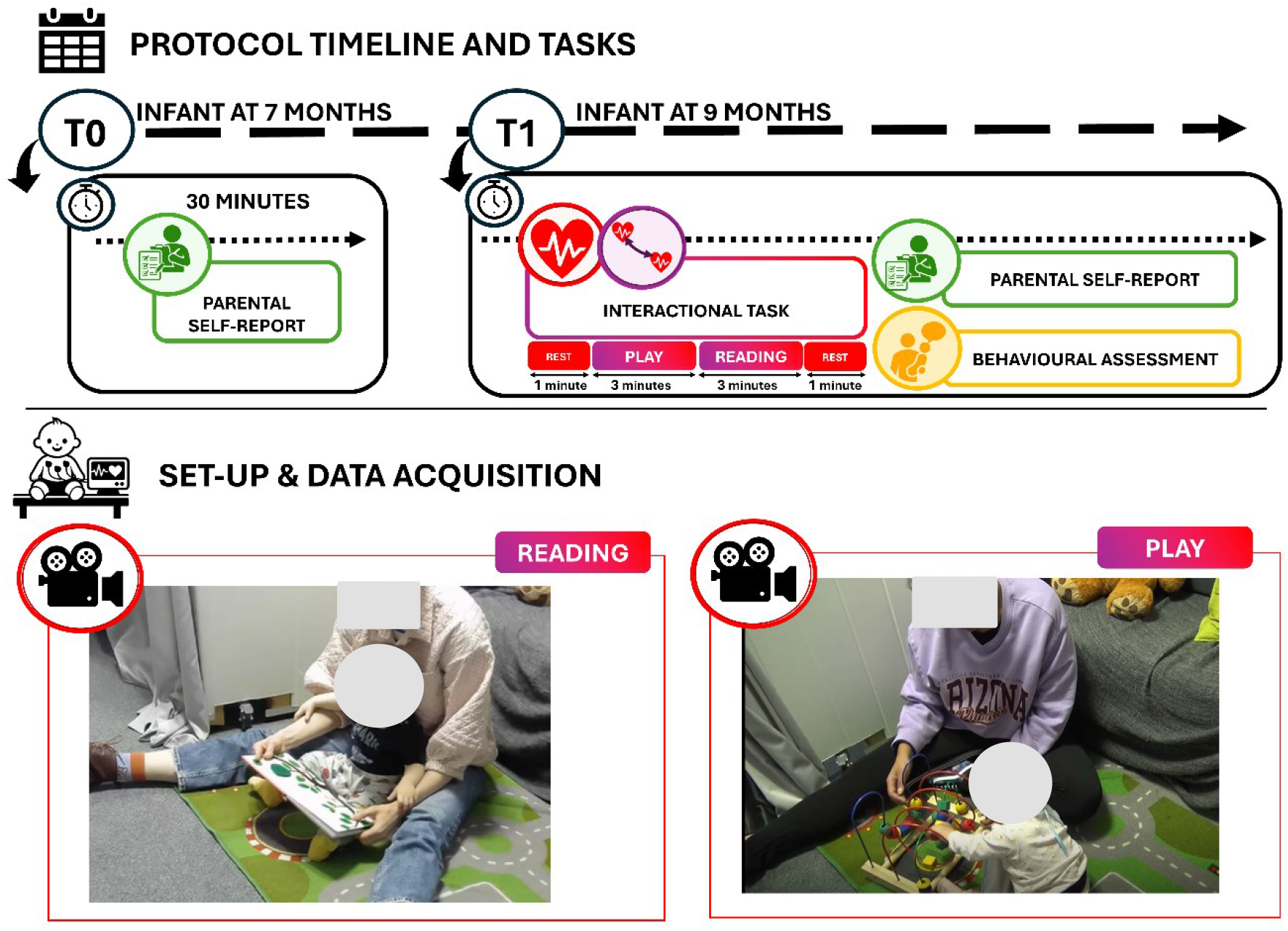
Study timeline and experimental design. Objectives are color-coded as follows: parental self-report (green), experimenter-administered behavioral assessment (yellow), autonomic measures (red), and autonomic synchrony (purple).

### Protocol description

A two-month intervention program was implemented when preterm infants reached 7 months of corrected age. Families were instructed to engage in shared book reading for at least 10 minutes per day, a minimum of four times per week, over an 8-week period. They were provided initial training regarding the strategies of shared book reading via personal meeting and video-tutorial and were given reading kits composed of 6 books selected to engage that age range by the Swiss Association “*Nés pour lire*”. The set of books was designed to elicit different types of parent–child interaction. Two titles were explicitly interactive, encouraging engagement between the parent, child, and book (On se cache, Nathan, 2013; Doudou lapin, Chétaud, 2016). One wordless book invited adults to create their own narratives and describe the illustrations (Bonne nuit, petit ourson, Canonica, 2019), whereas another prompted vocalizations through sound-based play (Un livre qui fait des sons, Tullet, 2017). The set also included a tactile book (Regarde mes amis, Weninger, 2019) and a book focused on facial emotion expressions (Mon imagier des émotions, Bost, 2017).

During the training session, parents were encouraged to adopt an interactive and emotionally responsive reading style, emphasizing shared attention, turn-taking, sensitivity to infant vocalizations and gestures, and the active co-construction of the interaction. They were specifically instructed to follow the infant’s attentional focus, verbally enrich the infant’s reactions, and maintain an emotionally engaging interaction throughout the reading activity. The tutorials highlighted the importance of reciprocal parent–infant exchanges and encouraged parents to adapt the pace and style of reading to the infant’s behavioral and emotional cues.

To assess the efficacy of the intervention, a parallel intervention was proposed to a second group of preterm born infants to create an active control group.

The shared play kit consisted of a set of colorful building blocks. Families were directed to spend the same amount of time in a spatial/building activity; provided equivalent training in how to mediate the infant-object interaction and techniques for shared play.

The shared play intervention was based on four core principles: (1) encouraging the infant to actively participate in the construction activity together with the parent; (2) observing and responding to the infant’s gestures, vocalizations, and behavioral cues; (3) building on the infant’s spontaneous interests and preferred forms of play; and (4) enriching the interaction through additional gestures, verbal comments, and playful proposals. Parents were encouraged to maintain an emotionally available and reciprocal interactional style throughout the activity, similarly to the shared reading condition.

Protocol adherence was monitored through biweekly video recordings of parent–infant interactions, which were reviewed by senior researchers who provided targeted feedback to ensure fidelity to key principles of dialogic reading, including turn-taking, sensitivity to infant cues, and reciprocal regulation. In addition, parents’ questions were addressed throughout the intervention, and adherence to the prescribed frequency and duration of reading sessions was monitored and reinforced. As a control for potential confounds, families completed a weekly questionnaire assessing the frequency of shared parent–child activities (e.g., outdoor outings, singing, dancing). Each activity was rated on a 7-point scale (0–6), and a composite score was computed across items. No significant group differences were observed (Kruskal–Wallis, *p* = 0.199).

#### Interactional Task

After the intervention, dyads were assessed at 9 months of corrected age for the preterm groups (PI and PC) and at 9 months of age for the term group (T). Each dyad participated in a structured sequence of interaction tasks, including an initial baseline, free play, and shared reading sessions presented in randomized order using novel toys or books, followed by a final baseline.

Throughout the session, physiological signals were continuously recorded from both mother and infant. Electrocardiographic (ECG) activity was simultaneously acquired using a BIOPAC MP160 system (BIOPAC Systems Inc., Goleta, CA) at a sampling rate of 1000 Hz. Signals were collected via wearable, portable sensors placed on the chest of both participants following standard pediatric and adult electrode configurations, allowing for synchronous recording of cardiac activity within each dyad.

All interactions took place in a controlled, infant-friendly laboratory environment with a soft, carpeted floor. To minimize external influences on behavior, the experimenter remained out of sight, and all sessions were audio- and video-recorded for subsequent analysis.

#### Behavioural assessment

Behavioural assessments were conducted using the Early Social Communication Scales (ESCS, Murray et al. 1990 (91)) to evaluate early social-communication abilities. All sessions were video-recorded for offline coding. Early social-communication behaviors were assessed using the Early Social Communication Scales (ESCS), focusing on three domains: Joint Attention (JA), Behavioral Requests (BR), and Social Interaction (SI). Behaviors were coded as either child-initiated or in response to tester prompts. BR captured the use of nonverbal behaviors to request assistance or objects, whereas SI reflected engagement in reciprocal, affectively positive interactions. JA was further subdivided into Initiating Joint Attention (IJA), indexing the use of eye contact, pointing, and showing to direct others’ attention, and Responding to Joint Attention (RJA), assessing the child’s ability to follow gaze and pointing cues.

#### Parental-self report

Before and after the intervention, parents completed validated questionnaires assessing psychological well-being, parent–infant relationship characteristics, and environmental factors. These included the Relationship Scale Questionnaire (RSQ, Griffin et Bartholomew 1994 (92); version validated by Guédeney, Fermanian et Bifulco, 2010 (93)), the Parenting Stress Index (PSI, Abidin et al 1995 (94)), and the Edinburgh Postpartum Depression Scale (EPDS, Murray et al. 1990 (95)). In addition, parents provided information on socio-economic status (SES) and completed a questionnaire assessing the weekly frequency of shared parent–infant activities (e.g., outdoor outings, play, and other interactive activities).

The EPDS was used to screen for symptoms of major postpartum depression (96).

The PSI assessed sources of stress related to parenthood, with a particular focus on stress experienced in the parental role (97).

The RSQ measured attachment style based on Bowlby’s theoretical framework, which links attachment patterns to internal representations of the self and others shaped by early relational experiences (93, 98). Within this framework, a secure attachment style reflects a positive model of both self and others, characterized by self-esteem and the expectation that others are available and responsive. In contrast, a fearful attachment style reflects negative models of both self and others, involving low self-worth and the perception of others as unreliable or unkind (93).

### ECG Data processing

#### Infant Autonomic Activation

R-peaks were detected, and artefacts and ectopic beats were corrected using Kubios HRV 2.2. Further analyses were performed in MATLAB 2021b®.

Interbeat interval (IBI) series were then segmented into 1-minute epochs for each task, excluding transition periods by selecting the second minute of shared tasks (Play and Reading). Vagal activation was quantified in the time domain using RMSSD, in the frequency domain using high-frequency power (HFpower) in the band [0.2–1.5 Hz] adapted to infants as reported (99), and in the nonlinear domain using SD1 (100). Cardiovascular complexity was assessed via Sample Entropy and the Complexity Index (CI), computed from Fuzzy Entropy across coarse-grained (101) across the first three scales, with CI defined as the area under the entropy curve.

#### Physiological Synchrony

IBI series during the 3-minutes interactional tasks (Play and Reading) were used to assess synchrony. To isolate respiration-independent cardiac activity, a respiratory signal (ECG-DR) was derived from the ECG using the slope-range method (102) and resampled at 4 Hz after outlier correction, performed as (103). The Orthogonal Subspace Projection (OSP) method (73) was then applied to decompose HRV into respiratory and residual components.

Both components have been used to assess synchrony separately, at respiratory and residual level respectively, after z-score normalization. This decomposition was essential given respiration variability during speech in the reading task, which would affect frequency-based autonomic and synchrony metrics. Cardiac-level synchrony using IBI has been previously demonstrated (104, 105).

#### Cross-Recurrence Quantification Analysis (CRQA)

CRQA (106) was applied to the respiratory and residual components to quantify dyadic physiological coupling (107, 108). The recurrence rate was fixed at 2%, according to (107, 108) for comparability across tasks and populations, with the time delay (τ) determined as the first local minimum of the auto-mutual information function and the embedding dimension (m) estimated using the False Nearest Neighbors (FNN) method (109). From the CRQA, the Rate of determinism (DET) was used as metric of synchrony (108), defined as the sum of diagonally adjacent recurrent points, higher values represent stronger coupling.

### Statistical Modelling

#### Missing data handling

73.12% percent of participants had at least one missing value in their demographic data, with the total fraction missing data in demographic variables 12.90%. In order to preserve observed data that would be removed in the presence of missing values, missing demographic data were imputed using classification and regression trees (CART). In total, 10 imputed data sets were created this way, and imputed values then averaged to create a final imputed data set.

#### Propensity score modelling for baseline confounding

To control for baseline confounding between the three study groups, we applied a propensity score approach (110–112). First, we fitted a multinomial logistic regression model with group as the outcome (terms as the reference level), and sex, SES, maternal age, and corrected infant age at 9 months as predictors. Demographic variables strongly distinguishing preterm and term infants (gestational age, birth weight, head circumference, Apgar scores) were excluded from this model. Two sets of fitted probabilities (i.e., propensity scores) were extracted from this model, one for the probability of being a pre-term intervention infant versus term infant, and another for the probability of being a pre-term control infant versus term infant. All subsequent analyses were then controlled for these two propensity scores, which ensured that group differences controlled for each participant’s baseline probability of belonging to their group. The advantage of this approach was that all confounder information was efficiently compressed into the propensity scores, without the need to do extensive further covariate adjustment in the target analyses.

#### Preterm Baseline Comparability

To assess homogeneity between the preterm groups, demographic variables not included in the propensity score model (corrected age at testing T0, Apgar scores, gestational age, birth weight, head circumference, weight and height percentiles) and T0 questionnaire scores were compared. For each variable, an ordinary linear regression was fitted with Group as a between-subject factor and propensity scores as covariates.

ANOVA Type II F-tests were used to evaluate group differences.

#### Questionnaire and Behavioral Outcomes at T1

Questionnaire subscales and behavioral scores at T1 were analyzed using ordinary regression with Group as a between-subject factor and propensity scores as covariates. Linear models were fitted separately for each outcome. ANOVA Type II F-tests were used to evaluate group differences, with degrees of freedom based on the residuals of the linear model. Significant effects were followed by Tukey-corrected post-hoc pairwise comparisons.

#### Autonomic Data

HRV indices were analyzed using linear mixed-effects models to account for the 3×4 design (Group × Task: Initial Baseline, Play, Reading, Final Baseline). Random intercepts were included for participants, and propensity scores were included as covariates.

ANOVA Type II Wald F-tests were used to evaluate group and task related differences, with degrees of freedom estimated using Satterthwaite’s approximation. Significant effects were followed by Tukey-corrected post-hoc pairwise comparisons. Estimated marginal means and Tukey-corrected post-hoc comparisons were computed to examine group differences across phases and phase differences within groups.

#### Cardiac Synchrony Assessment

Parent–infant cardiac synchrony was quantified using a surrogate data approach. Surrogate dyads were generated by pairing each parent’s signal with all non-corresponding children across groups and phases, excluding the true dyad. Synchrony parameters were extracted for all surrogate pairings, producing null distributions matched in duration and preprocessing. Real dyad synchrony metrics were compared against the surrogate distributions using two-sample Kolmogorov–Smirnov tests to assess whether observed coupling exceeded chance levels, as in (113).

## Author contributions

M.F., and D.G. designed research; L.L., F.B.-M, and C.B.-T., collected the data; L.L. and M.N. performed data analysis; L.L. an B.M. performed statistical modelling; L.L., M.F., and M.N. wrote the manuscript; P.H., E.G., D.G., S.D., L.C., M.N. and E.P.S. provided critical revisions and contributed to the interpretation of the data; all authors revised and approved the final version of the manuscript.

## Acknowledgments

We thank all the families who took part in this research project. Additional acknowledgment to the students and interns that assisted with subjects’ recruitment and data collection: Sonia Attianese, Magali Cynthia Uldry, Diana Guillen Vargas and Sabrina Bärtschi, and to the Swiss Centre for Affective Sciences.

