## Supplementary Tables for "Shared book reading promotes experience-dependent autonomic synchrony in parent–preterm infant dyads"

**Table S1. Preterm demographic data comparison**. Results from linear mixed-effects models fitted to neonatal demographic information. The first column lists model terms. Group denotes the two preterm infant groups (Preterm Intervention, Preterm Control). *propPI* and *propPC* indicate propensity scores, modeled as described in Materials and Methods.

| **Age at T0 corrected** | | | | |
|---|---|---|---|---|
| | *df* | *F-value* | *p-value* | η²partial |
| *Group* | 1 | 0,152 | 0,6993 | 5,08E-03 |
| *propPI* | 1 | 6,497 | 0,0150 | 2,88E-01 |
| *propPC* | 1 | 0,374 | 0,5444 | 9,75E-03 |
| *Residuals* | 38 | | | |
| **Age at T0 uncorrected** | | | | |
| | *df* | *F-value* | *p-value* | *η²partial* |
| *Group* | 1 | 0,220 | 0,6421 | 1,26E-02 |
| *propPI* | 1 | 1,611 | 0,2121 | 1,85E-01 |
| *propPC* | 1 | 2,055 | 0,1599 | 5,13E-02 |
| *Residuals* | 38 | | | |
| **APGAR 1m** | | | | |
| | *df* | *F-value* | *p-value* | η²partial |
| *Group* | 1 | 1,460 | 0,2328 | 2,90E-02 |
| *propPI* | 1 | 0,146 | 0,7045 | 7,62E-03 |
| *propPC* | 1 | 0,009 | 0,9262 | 1,81E-04 |
| *Residuals* | 48 | | | |
| **APGAR 5m** | | | | |
| | *df* | *F-value* | *p-value* | *η²partial* |
| *Group* | 1 | 1,972 | 0,1667 | 4,12E-02 |
| *propPI* | 1 | 0,130 | 0,7195 | 1,42E-02 |
| *propPC* | 1 | 0,132 | 0,7181 | 2,74E-03 |
| *Residuals* | 48 | | | |
| **APGAR 10m** | | | | |
| | *df* | *F-value* | *p-value* | *η²partial* |
| *Group* | 1 | 0,957 | 0,3329 | 3,04E-02 |
| *propPI* | 1 | 0,731 | 0,3967 | 1,15E-02 |
| *propPC* | 1 | 0,206 | 0,6523 | 4,27E-03 |
| *Residuals* | 48 | | | |
| **GA (days)** | | | | |
| | *df* | *F-value* | *p-value* | *η²partial* |
| *Group* | 1 | 0,245 | 0,6226 | 2,90E-04 |
| *propPI* | 1 | 4,320 | 0,0428 | 4,98E-02 |
| *propPC* | 1 | 1,709 | 0,1972 | 3,30E-02 |
| *Residuals* | 50 | | | |

| | df | F-value | p-value | η²partial |
|---|---|---|---|---|
| **Weight at birth (gr)** | | | | |
| *Group* | 1 | 1,388 | 0,2444 | 4,62E-02 |
| *propPI* | 1 | 1,118 | 0,2955 | 1,74E-03 |
| *propPC* | 1 | 3,751 | 0,0584 | 6,98E-02 |
| *Residuals* | 50 | | | |
| **Cranial Perimeter (cm)** | | | | |
| | *df* | *F-value* | *p-value* | *η²partial* |
| *Group* | 1 | 0,841 | 0,3634 | 3,47E-02 |
| *propPI* | 1 | 2,406 | 0,1272 | 2,90E-03 |
| *propPC* | 1 | 3,668 | 0,0612 | 6,84E-02 |
| *Residuals* | 50 | | | |
| **Percentile w** | | | | |
| | *df* | *F-value* | *p-value* | *η²partial* |
| *Group* | 1 | 0,542 | 0,4649 | 1,46E-02 |
| *propPI* | 1 | 0,120 | 0,7308 | 2,84E-05 |
| *propPC* | 1 | 0,321 | 0,5733 | 6,39E-03 |
| *Residuals* | 50 | | | |
| **Percentile p** | | | | |
| | *df* | *F-value* | *p-value* | *η²partial* |
| *Group* | 1 | 1,936 | 0,1702 | 4,35E-02 |
| *propPI* | 1 | 0,534 | 0,4684 | 1,67E-02 |
| *propPC* | 1 | 0,003 | 0,9532 | 6,95E-05 |
| *Residuals* | 50 | | | |

**Table S2. Preterm infants' parental report before the intervention (T0) comparison.** Results from linear mixed-effects models fitted to parental report scores at first. The first column lists model terms. *Group* denotes the two preterm infant groups (Preterm Intervention, Preterm Control). *propPI* and *propPC* indicate propensity scores, modeled as described in Materials and Methods.

| | | | | |
|---|---|---|---|---|
| **PSI** | | | | |
| **Total stress** | | | | |
| | *df* | *F-value* | *p-value* | η²partial |
| *Group* | 1 | 0,343 | 0,5651 | 0,015 |
| *propPI* | 1 | 0,034 | 0,8550 | 0,015 |
| *propPC* | 1 | 0,145 | 0,7073 | 0,008 |
| *Residuals* | 19 | | | |
| **Parental distress** | | | | |
| | *df* | *F-value* | *p-value* | η²partial |
| *Group* | 1 | 0,004 | 0,9484 | 0,001 |
| *propPI* | 1 | 0,788 | 0,3858 | 0,049 |
| *propPC* | 1 | 0,033 | 0,8575 | 0,002 |
| *Residuals* | 19 | | | |
| **Parent-child disfunction** | | | | |
| | *df* | *F-value* | *p-value* | η²partial |
| *Group* | 1 | 0,461 | 0,5053 | 0,023 |
| *propPI* | 1 | 0,043 | 0,8376 | 0,011 |
| *propPC* | 1 | 0,066 | 0,7994 | 0,003 |
| *Residuals* | 19 | | | |
| **Difficult child** | | | | |
| | *df* | *F-value* | *p-value* | η²partial |
| *Group* | 1 | 1,047 | 0,3191 | 0,031 |
| *propPI* | 1 | 0,903 | 0,3538 | 0,005 |
| *propPC* | 1 | 1,354 | 0,2590 | 0,067 |
| *Residuals* | 19 | | | |
| **RSQ** | | | | |
| **Total** | | | | |
| | *df* | *F-value* | *p-value* | η²partial |
| *Group* | 1 | 0,001 | 0,9788 | 0,002 |
| *propPI* | 1 | 0,553 | 0,4652 | 0,006 |
| *propPC* | 1 | 0,485 | 0,4938 | 0,023 |
| *Residuals* | 21 | | | |
| **F1** | | | | |
| | *df* | *F-value* | *p-value* | η²partial |
| *Group* | 1 | 0,706 | 0,4102 | 0,035 |
| *propPI* | 1 | 0,832 | 0,3721 | 0,042 |
| *propPC* | 1 | 0,100 | 0,7555 | 0,005 |
| *Residuals* | 21 | | | |

| F2 | | | | |
|---|---|---|---|---|
| | *df* | *F-value* | *p-value* | η²partial |
| *Group* | 1 | 0,107 | 0,7473 | 0,011 |
| *propPI* | 1 | 0,970 | 0,3358 | 0,020 |
| *propPC* | 1 | 0,541 | 0,4701 | 0,025 |
| *Residuals* | 21 | | | |
| **F3** | | | | |
| | *df* | *F-value* | *p-value* | η²partial |
| *Group* | 1 | 0,671 | 0,4219 | 0,053 |
| *propPI* | 1 | 0,077 | 0,7845 | 0,004 |
| *propPC* | 1 | 0,508 | 0,4839 | 0,024 |
| *Residuals* | 21 | | | |

| EPDS | | | | |
|---|---|---|---|---|
| | *df* | *F-value* | *p-value* | η²partial |
| *Group* | 1 | 0,036 | 0,8510 | 0,001 |
| *propPI* | 1 | 0,191 | 0,6671 | 0,011 |
| *propPC* | 1 | 0,027 | 0,8721 | 0,001 |
| *Residuals* | 19 | | | |

**Table S3. Cardiac Synchrony assessment.** Results from comparison of real synchrony against surrogate dyads.

| Respiratory component | | | |
|---|---|---|---|
| **Group** | **Task** | **p-value** | **K-stat** |
| Term | Play | 0.4048 | 0.121 |
| | Reading | 0.7279 | 0.074 |
| Preterm Intervention | Play | 0.9863 | 0.020 |
| | Reading | 0.0024 | 0.446 |
| Preterm Control | Play | 0.8594 | 0.064 |
| | Reading | 0.2583 | 0.187 |

| Residual component | | | |
|---|---|---|---|
| **Group** | **Task** | **p-value** | **K-stat** |
| Term | Play | 0.0198 | 0.253 |
| | Reading | 0.0050 | 0.305 |
| Preterm Intervention | Play | 0.9074 | 0.055 |
| | Reading | 0.9889 | 0.019 |
| Preterm Control | Play | 0.6651 | 0.106 |
| | Reading | 0.6860 | 0.099 |

**Table S4. Infant Autonomic assessment at test, comparison after the intervention (T1).** Results from linear mixed-effects models fitted to autonomic scores. The first column lists model terms. *Group* denotes the three infant groups (Term, Preterm Intervention, Preterm Control). *Task* denotes the four protocol activities (Initial Baseline, Play, Reading, Final Baseline). *propPI* and *propPC* indicate propensity scores, modeled as described in Materials and Methods.

| | | | | | |
|---|---|---|---|---|---|
| **Parasympathetic indices** | | | | | |
| **RMSSD** | | | | | |
| | *NumDF* | *DenDF* | *F-value* | *p-value* | η²partial |
| *Group* | 2 | 63,991 | 0,205 | 0,8149 | 0,007 |
| *Task* | 3 | 193,164 | 3,489 | 0,0168 | 0,059 |
| *propPI* | 1 | 63,832 | 0,206 | 0,6518 | 0,003 |
| *propPC* | 1 | 63,998 | 0,095 | 0,7595 | 0,001 |
| *Group:Task* | 6 | 193,208 | 1,360 | 0,2325 | 0,041 |
| **HF** | | | | | |
| | *NumDF* | *DenDF* | *F-value* | *p-value* | η²partial |
| *Group* | 2 | 64,394 | 0,122 | 0,8853 | 0,004 |
| *Task* | 3 | 193,738 | 3,143 | 0,0264 | 0,054 |
| *propPI* | 1 | 64,094 | 0,180 | 0,6729 | 0,003 |
| *propPC* | 1 | 64,409 | 0,127 | 0,7225 | 0,002 |
| *Group:Task* | 6 | 193,819 | 1,927 | 0,0782 | 0,056 |
| **SD1** | | | | | |
| | *NumDF* | *DenDF* | *F-value* | *p-value* | η²partial |
| *Group* | 2 | 63,990 | 0,205 | 0,8149 | 0,007 |
| *Task* | 3 | 193,163 | 3,489 | 0,0168 | 0,059 |
| *propPI* | 1 | 63,832 | 0,204 | 0,6530 | 0,003 |
| *propPC* | 1 | 63,998 | 0,094 | 0,7607 | 0,001 |
| *Group:Task* | 6 | 193,207 | 1,360 | 0,2327 | 0,041 |
| **Entropy indeces** | | | | | |
| **Sample entopy** | | | | | |
| | *NumDF* | *DenDF* | *F-value* | *p-value* | η²partial |
| *Group* | 2 | 65,014 | 4,827 | 0,0111 | 0,128 |
| *Task* | 3 | 194,588 | 3,435 | 0,0180 | 0,070 |
| *propPI* | 1 | 64,537 | 0,015 | 0,9020 | 0,000 |
| *propPC* | 1 | 65,040 | 0,812 | 0,3710 | 0,012 |
| *Group:Task* | 6 | 194,714 | 2,040 | 0,0621 | 0,059 |
| **CI** | | | | | |
| | *NumDF* | *DenDF* | *F-value* | *p-value* | η²partial |
| *Group* | 2 | 64,789 | 3,266 | 0,0445 | 0,091 |
| *Task* | 3 | 194,375 | 1,372 | 0,2527 | 0,036 |
| *propPI* | 1 | 64,309 | 0,001 | 0,9774 | 0,000 |
| *propPC* | 1 | 64,815 | 0,128 | 0,7216 | 0,002 |
| *Group:Task* | 6 | 194,502 | 2,178 | 0,0467 | 0,063 |

**Table S5. Post-hoc analysis on RMSSD**. The Fix term column reports the following abbreviations T = Term, PI = Preterm Intervention, PC = Preterm Control.

#### RMSSD

| | Fixed term | contrast | Mean Diff | SE | df | t-ratio | p-value | 95% CI | Cohen's d | 95% CI (d) |
|---|---|---|---|---|---|---|---|---|---|---|
| Between-group | Initial Baseline | Control - Intervention | -2,66E-03 | 1,86E-03 | 104,703 | -1,432 | 0,3283 | [-7,08E-03;1,76E-03] | -0,856 | [-2,04E+00;3,33E-01] |
| | | Control - Term | -8,46E-04 | 1,87E-03 | 90,194 | -0,453 | 0,8933 | [-5,29E-03;3,60E-03] | -0,272 | [-1,46E+00;9,21E-01] |
| | | Intervention - Term | 1,81E-03 | 1,98E-03 | 92,12 | 0,918 | 0,6307 | [-2,90E-03;6,53E-03] | 0,584 | [-6,81E-01;1,85E+00] |
| | Play | Control - Intervention | -3,54E-04 | 1,86E-03 | 104,703 | -0,191 | 0,9802 | [-4,77E-03;4,06E-03] | -0,114 | [-1,30E+00;1,07E+00] |
| | | Control - Term | -5,41E-04 | 1,87E-03 | 90,194 | -0,29 | 0,9547 | [-4,99E-03;3,91E-03] | -0,174 | [-1,37E+00;1,02E+00] |
| | | Intervention - Term | -1,87E-04 | 1,98E-03 | 92,12 | -0,095 | 0,9951 | [-4,90E-03;4,52E-03] | -0,06 | [-1,32E+00;1,20E+00] |
| | Reading | Control - Intervention | 4,07E-04 | 1,86E-03 | 104,703 | 0,219 | 0,9739 | [-4,01E-03;4,82E-03] | 0,131 | [-1,06E+00;1,32E+00] |
| | | Control - Term | 1,25E-03 | 1,87E-03 | 90,194 | 0,672 | 0,7805 | [-3,20E-03;5,70E-03] | 0,403 | [-7,90E-01;1,60E+00] |
| | | Intervention - Term | 8,47E-04 | 1,98E-03 | 92,12 | 0,428 | 0,9039 | [-3,86E-03;5,56E-03] | 0,273 | [-9,91E-01;1,54E+00] |
| | Final Baseline | Control - Intervention | -7,95E-04 | 1,89E-03 | 111,763 | -0,42 | 0,9075 | [-5,30E-03;3,71E-03] | -0,256 | [-1,47E+00;9,54E-01] |
| | | Control - Term | 1,07E-03 | 1,88E-03 | 93,074 | 0,568 | 0,8375 | [-3,42E-03;5,56E-03] | 0,344 | [-8,59E-01;1,55E+00] |
| | | Intervention - Term | 1,86E-03 | 2,02E-03 | 98,761 | 0,924 | 0,6263 | [-2,94E-03;6,67E-03] | 0,6 | [-6,89E-01;1,89E+00] |
| Within-group | PC | Initial Baseline - Play | 9,35E-04 | 9,59E-04 | 193,005 | 0,975 | 0,7641 | [-1,55E-03;3,42E-03] | 0,301 | [-3,12E-01;9,14E-01] |
| | | Initial Baseline - Reading | -5,14E-04 | 9,59E-04 | 193,005 | -0,536 | 0,9502 | [-3,00E-03;1,97E-03] | -0,165 | [-7,78E-01;4,47E-01] |
| | | Initial Baseline - Final Baseline | -7,34E-04 | 9,75E-04 | 193,377 | -0,753 | 0,8752 | [-3,26E-03;1,79E-03] | -0,236 | [-8,59E-01;3,86E-01] |
| | | Play - Reading | -1,45E-03 | 9,59E-04 | 193,005 | -1,51 | 0,4333 | [-3,93E-03;1,04E-03] | -0,466 | [-1,08E+00;1,48E-01] |
| | | Play – Final Baseline | -1,67E-03 | 9,75E-04 | 193,377 | -1,712 | 0,32 | [-4,20E-03;8,57E-04] | -0,537 | [-1,16E+00;8,71E-02] |
| | | Reading - Final Baseline | -2,20E-04 | 9,75E-04 | 193,377 | -0,226 | 0,9959 | [-2,75E-03;2,31E-03] | -0,071 | [-6,93E-01;5,51E-01] |
| | PI | Initial Baseline - Play | 3,24E-03 | 1,10E-03 | 193,005 | 2,949 | 0,0186 | [3,93E-04;6,09E-03] | 1,043 | [3,35E-01;1,75E+00] |
| | | Initial Baseline - Reading | 2,55E-03 | 1,10E-03 | 193,005 | 2,323 | 0,0964 | [-2,95E-04;5,40E-03] | 0,821 | [1,16E-01;1,53E+00] |
| | | Initial Baseline - Final Baseline | 1,13E-03 | 1,15E-03 | 194,058 | 0,984 | 0,7586 | [-1,85E-03;4,11E-03] | 0,364 | [-3,70E-01;1,10E+00] |
| | | Play - Reading | -6,88E-04 | 1,10E-03 | 193,005 | -0,626 | 0,9236 | [-3,54E-03;2,16E-03] | -0,221 | [-9,23E-01;4,80E-01] |
| | | Play – Final Baseline | -2,11E-03 | 1,15E-03 | 194,058 | -1,837 | 0,2591 | [-5,09E-03;8,66E-04] | -0,679 | [-1,41E+00;5,67E-02] |
| | | Reading - Final Baseline | -1,42E-03 | 1,15E-03 | 194,058 | -1,238 | 0,6034 | [-4,40E-03;1,55E-03] | -0,458 | [-1,19E+00;2,77E-01] |
| | T | Initial Baseline - Play | 1,24E-03 | 7,77E-04 | 193,005 | 1,595 | 0,3841 | [-7,75E-04;3,25E-03] | 0,399 | [-9,90E-02;8,96E-01] |
| | | Initial Baseline - Reading | 1,59E-03 | 7,77E-04 | 193,005 | 2,04 | 0,1769 | [-4,28E-04;3,60E-03] | 0,51 | [1,16E-02;1,01E+00] |

| | | | | | | | | | |
|---|---|---|---|---|---|---|---|---|---|
| | Initial Baseline - Final Baseline | 1,18E-03 | 7,94E-04 | 193,495 | 1,488 | 0,4468 | [-8,76E-04;3,24E-03] | 0,38 | [-1,28E-01;8,88E-01] |
| | Play - Reading | 3,46E-04 | 7,77E-04 | 193,005 | 0,446 | 0,9704 | [-1,67E-03;2,36E-03] | 0,111 | [-3,85E-01;6,08E-01] |
| | Play - Final Baseline | -5,82E-05 | 7,94E-04 | 193,495 | -0,073 | 0,9999 | [-2,12E-03;2,00E-03] | -0,019 | [-5,26E-01;4,88E-01] |
| | Reading - Final Baseline | -4,05E-04 | 7,94E-04 | 193,495 | -0,51 | 0,9567 | [-2,46E-03;1,65E-03] | -0,13 | [-6,37E-01;3,77E-01] |

**Table S6. Post-hoc analysis on HF**. The Fix term column reports the following abbreviations T = Term, PI = Preterm Intervention, PC = Preterm Control.

## HF

| | Fixed term | contrast | Mean Diff | SE | df | t-ratio | p-value | 95% CI | Cohen's d | 95% CI (d) |
|---|---|---|---|---|---|---|---|---|---|---|
| Between-group | Initial Baseline | Control - Intervention | -4,92E-05 | 2,77E-05 | 145,893 | -1,773 | 0,1822 | [-1,15E-04;1,65E-05] | -0,831 | [-1,76E+00;9,87E-02] |
| | | Control - Term | -2,64E-05 | 2,71E-05 | 117,667 | -0,973 | 0,5951 | [-9,07E-05;3,80E-05] | -0,446 | [-1,35E+00;4,61E-01] |
| | | Intervention - Term | 2,28E-05 | 2,88E-05 | 121,538 | 0,79 | 0,7096 | [-4,56E-05;9,12E-05] | 0,385 | [-5,79E-01;1,35E+00] |
| | Play | Control - Intervention | -3,85E-06 | 2,77E-05 | 145,893 | -0,139 | 0,9894 | [-6,95E-05;6,18E-05] | -0,065 | [-9,92E-01;8,61E-01] |
| | | Control - Term | -8,12E-06 | 2,71E-05 | 117,667 | -0,3 | 0,9517 | [-7,25E-05;5,62E-05] | -0,137 | [-1,04E+00;7,69E-01] |
| | | Intervention - Term | -4,27E-06 | 2,88E-05 | 121,538 | -0,148 | 0,988 | [-7,27E-05;6,41E-05] | -0,072 | [-1,04E+00;8,92E-01] |
| | Reading | Control - Intervention | 1,43E-05 | 2,77E-05 | 145,893 | 0,516 | 0,8636 | [-5,14E-05;8,00E-05] | 0,242 | [-6,85E-01;1,17E+00] |
| | | Control - Term | -3,65E-06 | 2,71E-05 | 117,667 | -0,135 | 0,9901 | [-6,80E-05;6,07E-05] | -0,062 | [-9,68E-01;8,44E-01] |
| | | Intervention - Term | -1,80E-05 | 2,88E-05 | 121,538 | -0,623 | 0,8078 | [-8,64E-05;5,05E-05] | -0,303 | [-1,27E+00;6,61E-01] |
| | Final Baseline | Control - Intervention | -4,03E-06 | 2,86E-05 | 157,506 | -0,141 | 0,9891 | [-7,17E-05;6,37E-05] | -0,068 | [-1,02E+00;8,87E-01] |
| | | Control - Term | 3,22E-05 | 2,75E-05 | 122,966 | 1,17 | 0,4731 | [-3,31E-05;9,75E-05] | 0,544 | [-3,77E-01;1,46E+00] |
| | | Intervention - Term | 3,62E-05 | 2,98E-05 | 133,53 | 1,216 | 0,4459 | [-3,44E-05;1,07E-04] | 0,612 | [-3,85E-01;1,61E+00] |
| Within-group | PC | Initial Baseline - Play | 1,06E-05 | 1,83E-05 | 193,017 | 0,579 | 0,9383 | [-3,68E-05;5,79E-05] | 0,179 | [-4,32E-01;7,89E-01] |
| | | Initial Baseline - Reading | -3,00E-06 | 1,83E-05 | 193,017 | -0,164 | 0,9984 | [-5,04E-05;4,44E-05] | -0,051 | [-6,61E-01;5,60E-01] |
| | | Initial Baseline - Final Baseline | -2,31E-05 | 1,86E-05 | 193,723 | -1,244 | 0,5997 | [-7,12E-05;2,50E-05] | -0,39 | [-1,01E+00;2,31E-01] |
| | | Play - Reading | -1,36E-05 | 1,83E-05 | 193,017 | -0,743 | 0,8795 | [-6,09E-05;3,38E-05] | -0,229 | [-8,40E-01;3,81E-01] |
| | | Play – Final Baseline | -3,37E-05 | 1,86E-05 | 193,723 | -1,814 | 0,2699 | [-8,18E-05;1,44E-05] | -0,569 | [-1,19E+00;5,34E-02] |
| | | Reading - Final Baseline | -2,01E-05 | 1,86E-05 | 193,723 | -1,082 | 0,7007 | [-6,82E-05;2,80E-05] | -0,339 | [-9,60E-01;2,81E-01] |
| | PI | Initial Baseline - Play | 5,59E-05 | 2,09E-05 | 193,017 | 2,67 | 0,0407 | [1,65E-06;1,10E-04] | 0,944 | [2,40E-01;1,65E+00] |
| | | Initial Baseline - Reading | 6,05E-05 | 2,09E-05 | 193,017 | 2,89 | 0,0222 | [6,24E-06;1,15E-04] | 1,022 | [3,17E-01;1,73E+00] |
| | | Initial Baseline - Final Baseline | 2,21E-05 | 2,19E-05 | 195,009 | 1,01 | 0,744 | [-3,46E-05;7,87E-05] | 0,373 | [-3,58E-01;1,10E+00] |
| | | Play - Reading | 4,59E-06 | 2,09E-05 | 193,017 | 0,219 | 0,9963 | [-4,97E-05;5,88E-05] | 0,078 | [-6,22E-01;7,77E-01] |
| | | Play – Final Baseline | -3,38E-05 | 2,19E-05 | 195,009 | -1,548 | 0,4107 | [-9,05E-05;2,28E-05] | -0,571 | [-1,30E+00;1,60E-01] |
| | | Reading - Final Baseline | -3,84E-05 | 2,19E-05 | 195,009 | -1,759 | 0,2966 | [-9,51E-05;1,82E-05] | -0,649 | [-1,38E+00;8,30E-02] |
| | T | Initial Baseline - Play | 2,88E-05 | 1,48E-05 | 193,017 | 1,948 | 0,2115 | [-9,52E-06;6,72E-05] | 0,487 | [-9,56E-03;9,84E-01] |
| | | Initial Baseline - Reading | 1,97E-05 | 1,48E-05 | 193,017 | 1,333 | 0,5429 | [-1,86E-05;5,81E-05] | 0,333 | [-1,62E-01;8,29E-01] |

| | | | | | | | | |
|---|---|---|---|---|---|---|---|---|
| Initial Baseline - Final Baseline | 3,55E-05 | 1,51E-05 | 193,947 | 2,349 | 0,0907 | [-3,66E-06;7,47E-05] | 0,6 | [9,17E-02;1,11E+00] |
| Play - Reading | -9,11E-06 | 1,48E-05 | 193,017 | -0,615 | 0,9272 | [-4,75E-05;2,93E-05] | -0,154 | [-6,49E-01;3,41E-01] |
| Play - Final Baseline | 6,66E-06 | 1,51E-05 | 193,947 | 0,44 | 0,9714 | [-3,25E-05;4,58E-05] | 0,112 | [-3,93E-01;6,18E-01] |
| Reading - Final Baseline | 1,58E-05 | 1,51E-05 | 193,947 | 1,043 | 0,7244 | [-2,34E-05;5,49E-05] | 0,266 | [-2,39E-01;7,72E-01] |

**Table S7. Post-hoc analysis on SD1**. The Fix term column reports the following abbreviations T = Term, PI = Preterm Intervention, PC = Preterm Control.

# SD1

| | Fixed term | contrast | Mean Diff | SE | df | t-ratio | p-value | 95% CI | Cohen's d | 95% CI (d) |
|---|---|---|---|---|---|---|---|---|---|---|
| Between-group | Initial Baseline | Control - Intervention | -1,89E-03 | 0,001 | 104,717 | -1,432 | 0,3283 | [-5,03E-03;1,25E-03] | -0,856 | [-2,04E+00;3,32E-01] |
| | | Control - Term | -6,01E-04 | 0,001 | 90,202 | -0,453 | 0,893 | [-3,76E-03;2,56E-03] | -0,272 | [-1,47E+00;9,20E-01] |
| | | Intervention - Term | 1,29E-03 | 0,001 | 92,129 | 0,917 | 0,631 | [-2,06E-03;4,63E-03] | 0,583 | [-6,81E-01;1,85E+00] |
| | Play | Control - Intervention | -2,52E-04 | 0,001 | 104,717 | -0,191 | 0,98 | [-3,39E-03;2,88E-03] | -0,114 | [-1,30E+00;1,07E+00] |
| | | Control - Term | -3,86E-04 | 0,001 | 90,202 | -0,291 | 0,9544 | [-3,55E-03;2,77E-03] | -0,175 | [-1,37E+00;1,02E+00] |
| | | Intervention - Term | -1,33E-04 | 0,001 | 92,129 | -0,095 | 0,995 | [-3,48E-03;3,21E-03] | -0,06 | [-1,32E+00;1,20E+00] |
| | Reading | Control - Intervention | 2,88E-04 | 0,001 | 104,717 | 0,218 | 0,9741 | [-2,85E-03;3,42E-03] | 0,13 | [-1,06E+00;1,32E+00] |
| | | Control - Term | 8,89E-04 | 0,001 | 90,202 | 0,671 | 0,7812 | [-2,27E-03;4,05E-03] | 0,403 | [-7,90E-01;1,60E+00] |
| | | Intervention - Term | 6,01E-04 | 0,001 | 92,129 | 0,428 | 0,9041 | [-2,75E-03;3,95E-03] | 0,272 | [-9,91E-01;1,54E+00] |
| | Final Baseline | Control - Intervention | -5,65E-04 | 0,001 | 111,778 | -0,42 | 0,9074 | [-3,76E-03;2,63E-03] | -0,256 | [-1,47E+00;9,53E-01] |
| | | Control - Term | 7,60E-04 | 0,001 | 93,083 | 0,568 | 0,8376 | [-2,43E-03;3,95E-03] | 0,344 | [-8,59E-01;1,55E+00] |
| | | Intervention - Term | 1,32E-03 | 0,001 | 98,772 | 0,924 | 0,6262 | [-2,09E-03;4,73E-03] | 0,6 | [-6,89E-01;1,89E+00] |
| Within-group | PC | Initial Baseline - Play | 6,65E-04 | 0,001 | 193,005 | 0,975 | 0,7636 | [-1,10E-03;2,43E-03] | 0,301 | [-3,12E-01;9,14E-01] |
| | | Initial Baseline - Reading | -3,64E-04 | 0,001 | 193,005 | -0,534 | 0,9506 | [-2,13E-03;1,40E-03] | -0,165 | [-7,78E-01;4,48E-01] |
| | | Initial Baseline - Final Baseline | -5,22E-04 | 0,001 | 193,377 | -0,753 | 0,8751 | [-2,32E-03;1,27E-03] | -0,236 | [-8,59E-01;3,86E-01] |
| | | Play - Reading | -1,03E-03 | 0,001 | 193,005 | -1,51 | 0,4336 | [-2,79E-03;7,37E-04] | -0,466 | [-1,08E+00;1,48E-01] |
| | | Play – Final Baseline | -1,19E-03 | 0,001 | 193,377 | -1,713 | 0,3195 | [-2,98E-03;6,08E-04] | -0,537 | [-1,16E+00;8,68E-02] |
| | | Reading - Final Baseline | -1,57E-04 | 0,001 | 193,377 | -0,227 | 0,9958 | [-1,95E-03;1,64E-03] | -0,071 | [-6,94E-01;5,51E-01] |
| | PI | Initial Baseline - Play | 2,30E-03 | 0,001 | 193,005 | 2,948 | 0,0187 | [2,78E-04;4,32E-03] | 1,042 | [3,35E-01;1,75E+00] |
| | | Initial Baseline - Reading | 1,81E-03 | 0,001 | 193,005 | 2,323 | 0,0964 | [-2,10E-04;3,84E-03] | 0,821 | [1,16E-01;1,53E+00] |
| | | Initial Baseline - Final Baseline | 8,03E-04 | 0,001 | 194,058 | 0,984 | 0,759 | [-1,31E-03;2,92E-03] | 0,363 | [-3,70E-01;1,10E+00] |
| | | Play - Reading | -4,88E-04 | 0,001 | 193,005 | -0,626 | 0,9238 | [-2,51E-03;1,53E-03] | -0,221 | [-9,23E-01;4,81E-01] |
| | | Play – Final Baseline | -1,50E-03 | 0,001 | 194,058 | -1,837 | 0,2591 | [-3,61E-03;6,15E-04] | -0,679 | [-1,41E+00;5,67E-02] |
| | | Reading - Final Baseline | -1,01E-03 | 0,001 | 194,058 | -1,239 | 0,6031 | [-3,13E-03;1,10E-03] | -0,458 | [-1,19E+00;2,77E-01] |
| | T | Initial Baseline - Play | 8,80E-04 | 0,001 | 193,005 | 1,594 | 0,3843 | [-5,51E-04;2,31E-03] | 0,399 | [-9,91E-02;8,96E-01] |
| | | Initial Baseline - Reading | 1,13E-03 | 0,001 | 193,005 | 2,04 | 0,1769 | [-3,04E-04;2,56E-03] | 0,51 | [1,16E-02;1,01E+00] |
| | | Initial Baseline - Final Baseline | 8,39E-04 | 0,001 | 193,496 | 1,488 | 0,4465 | [-6,22E-04;2,30E-03] | 0,38 | [-1,28E-01;8,88E-01] |
| | | Play - Reading | 2,46E-04 | 0,001 | 193,005 | 0,446 | 0,9703 | [-1,18E-03;1,68E-03] | 0,112 | [-3,85E-01;6,08E-01] |
| | | Play - Final Baseline | -4,09E-05 | 0,001 | 193,496 | -0,073 | 0,9999 | [-1,50E-03;1,42E-03] | -0,019 | [-5,26E-01;4,88E-01] |
| | | Reading - Final Baseline | -2,87E-04 | 0,001 | 193,496 | -0,509 | 0,9568 | [-1,75E-03;1,17E-03] | -0,13 | [-6,37E-01;3,77E-01] |

**Table S8. Post-hoc analysis on Sample Entropy**. The Fix term column report the following abbreviations T = Term, PI = Preterm Intervention, PC = Preterm Control.

### Sample Entropy

| | Fixed term | contrast | Mean Diff | SE | df | t-ratio | p-value | 95% CI | Cohen's d | 95% CI (d) |
|---|---|---|---|---|---|---|---|---|---|---|
| Between-group | Initial Baseline | Control - Intervention | -0,057 | 0,092 | 194,271 | -0,617 | 0,8109 | [-0,273;0,160] | -0,249 | [-1,044;0,547] |
| | | Control - Term | 0,04 | 0,087 | 154,977 | 0,464 | 0,888 | [-0,166;0,247] | 0,178 | [-0,579;0,935] |
| | | Intervention - Term | 0,097 | 0,093 | 160,869 | 1,042 | 0,5514 | [-0,123;0,317] | 0,427 | [-0,383;1,236] |
| | Play | Control - Intervention | 0 | 0,092 | 194,271 | 0,002 | 1 | [-0,216;0,216] | 0,001 | [-0,794;0,796] |
| | | Control - Term | 0,237 | 0,087 | 154,977 | 2,723 | 0,0196 | [0,031;0,443] | 1,044 | [0,281;1,806] |
| | | Intervention - Term | 0,237 | 0,093 | 160,869 | 2,548 | 0,0315 | [0,017;0,457] | 1,043 | [0,230;1,856] |
| | Reading | Control - Intervention | -0,232 | 0,092 | 194,271 | -2,535 | 0,0321 | [-0,448;-0,016] | -1,021 | [-1,821;-0,221] |
| | | Control - Term | 0,127 | 0,087 | 154,977 | 1,457 | 0,3145 | [-0,079;0,333] | 0,558 | [-0,200;1,317] |
| | | Intervention - Term | 0,359 | 0,093 | 160,869 | 3,859 | 0,0005 | [0,139;0,579] | 1,58 | [0,760;2,400] |
| | Final Baseline | Control - Intervention | -0,124 | 0,095 | 205,707 | -1,302 | 0,3959 | [-0,349;0,101] | -0,546 | [-1,374;0,283] |
| | | Control - Term | 0,066 | 0,089 | 161,926 | 0,739 | 0,7405 | [-0,145;0,276] | 0,289 | [-0,483;1,061] |
| | | Intervention - Term | 0,19 | 0,097 | 175,814 | 1,951 | 0,1276 | [-0,040;0,420] | 0,835 | [-0,013;1,682] |
| Within-group | PC | Initial_Baseline - Play | -0,197 | 0,07 | 193,048 | -2,801 | 0,0285 | [-0,378;-0,015] | -0,865 | [-1,478;-0,251] |
| | | Initial_Baseline - Reading | -0,117 | 0,07 | 193,048 | -1,674 | 0,34 | [-0,299;0,064] | -0,517 | [-1,127;0,094] |
| | | Initial_Baseline - Final Baseline | -0,071 | 0,071 | 194,165 | -0,992 | 0,7539 | [-0,255;0,114] | -0,311 | [-0,930;0,308] |
| | | Play - Reading | 0,079 | 0,07 | 193,048 | 1,127 | 0,6734 | [-0,103;0,261] | 0,348 | [-0,262;0,958] |
| | | Play - Final_Baseline | 0,126 | 0,071 | 194,165 | 1,767 | 0,2922 | [-0,059;0,310] | 0,554 | [-0,066;1,174] |
| | | Reading - Final Baseline | 0,047 | 0,071 | 194,165 | 0,657 | 0,9129 | [-0,138;0,231] | 0,206 | [-0,413;0,824] |
| | PI | Initial Baseline - Play | -0,14 | 0,08 | 193,048 | -1,74 | 0,3058 | [-0,348;0,068] | -0,615 | [-1,315;0,084] |
| | | Initial Baseline - Reading | -0,293 | 0,08 | 193,048 | -3,647 | 0,0019 | [-0,501;-0,085] | -1,289 | [-1,996;-0,583] |
| | | Initial Baseline - Final Baseline | -0,138 | 0,084 | 196,198 | -1,65 | 0,3533 | [-0,355;0,079] | -0,608 | [-1,337;0,121] |
| | | Play - Reading | -0,153 | 0,08 | 193,048 | -1,907 | 0,2285 | [-0,362;0,055] | -0,674 | [-1,374;0,026] |
| | | Play - Final_Baseline | 0,002 | 0,084 | 196,198 | 0,02 | 1 | [-0,215;0,219] | 0,007 | [-0,720;0,734] |
| | | Reading - Final Baseline | 0,155 | 0,084 | 196,198 | 1,85 | 0,2533 | [-0,062;0,372] | 0,682 | [-0,048;1,411] |
| | T | Initial Baseline - Play | 0 | 0,057 | 193,048 | 0,004 | 1 | [-0,147;0,148] | 0,001 | [-0,493;0,495] |
| | | Initial Baseline - Reading | -0,031 | 0,057 | 193,048 | -0,545 | 0,9477 | [-0,178;0,116] | -0,136 | [-0,630;0,357] |
| | | Initial Baseline - Final Baseline | -0,045 | 0,058 | 194,521 | -0,783 | 0,862 | [-0,196;0,105] | -0,2 | [-0,704;0,304] |
| | | Play - Reading | -0,031 | 0,057 | 193,048 | -0,549 | 0,9466 | [-0,179;0,116] | -0,137 | [-0,631;0,356] |
| | | Play - Final Baseline | -0,046 | 0,058 | 194,521 | -0,787 | 0,8601 | [-0,196;0,105] | -0,201 | [-0,705;0,303] |
| | | Reading - Final Baseline | -0,014 | 0,058 | 194,521 | -0,249 | 0,9946 | [-0,165;0,136] | -0,063 | [-0,567;0,440] |

**Table 9. Post-hoc analysis on Complexity Index (CI)**. The Fix term column report the following abbreviations T = Term, PI = Preterm Intervention, PC = Preterm Control.

## CI

| | Fixed term | contrast | Mean Diff | SE | df | t-ratio | p-value | 95% CI | Cohen's d | 95% CI (d) |
|---|---|---|---|---|---|---|---|---|---|---|
| Between-group | Initial Baseline | Control - Intervention | -0,052 | 0,101 | 195,253 | -0,517 | 0,8633 | [-0,290;0,186] | -0,208 | [-1,001;0,586] |
| | | Control - Term | 0,016 | 0,096 | 155,836 | 0,172 | 0,9838 | [-0,210;0,243] | 0,066 | [-0,689;0,820] |
| | | Intervention - Term | 0,069 | 0,102 | 161,762 | 0,67 | 0,7815 | [-0,174;0,311] | 0,273 | [-0,533;1,080] |
| | Play | Control - Intervention | 0,051 | 0,101 | 195,253 | 0,509 | 0,8672 | [-0,187;0,289] | 0,204 | [-0,589;0,997] |
| | | Control - Term | 0,212 | 0,096 | 155,836 | 2,209 | 0,0728 | [-0,015;0,439] | 0,844 | [0,086;1,602] |
| | | Intervention - Term | 0,16 | 0,102 | 161,762 | 1,567 | 0,2628 | [-0,082;0,403] | 0,64 | [-0,168;1,447] |
| | Reading | Control - Intervention | -0,261 | 0,101 | 195,253 | -2,591 | 0,0277 | [-0,499;-0,023] | -1,041 | [-1,839;-0,243] |
| | | Control - Term | 0,121 | 0,096 | 155,836 | 1,262 | 0,4187 | [-0,106;0,348] | 0,482 | [-0,273;1,238] |
| | | Intervention - Term | 0,382 | 0,102 | 161,762 | 3,732 | 0,0008 | [0,140;0,624] | 1,523 | [0,706;2,340] |
| | Final Baseline | Control - Intervention | -0,092 | 0,105 | 206,614 | -0,872 | 0,6584 | [-0,339;0,156] | -0,365 | [-1,190;0,461] |
| | | Control - Term | 0,096 | 0,098 | 162,796 | 0,986 | 0,5868 | [-0,135;0,328] | 0,384 | [-0,386;1,155] |
| | | Intervention - Term | 0,188 | 0,107 | 176,711 | 1,756 | 0,1876 | [-0,065;0,441] | 0,749 | [-0,095;1,594] |
| Within-group | PC | Initial Baseline - Play | -0,175 | 0,077 | 193,048 | -2,254 | 0,1127 | [-0,375;0,026] | -0,696 | [-1,308;-0,084] |
| | | Initial Baseline - Reading | -0,084 | 0,077 | 193,048 | -1,086 | 0,6984 | [-0,285;0,117] | -0,335 | [-0,945;0,275] |
| | | Initial Baseline - Final Baseline | -0,074 | 0,079 | 194,176 | -0,939 | 0,7839 | [-0,277;0,130] | -0,294 | [-0,913;0,325] |
| | | Play - Reading | 0,09 | 0,077 | 193,048 | 1,168 | 0,6479 | [-0,110;0,291] | 0,36 | [-0,249;0,970] |
| | | Play – Final Baseline | 0,101 | 0,079 | 194,176 | 1,282 | 0,5758 | [-0,103;0,304] | 0,401 | [-0,218;1,021] |
| | | Reading - Final Baseline | 0,01 | 0,079 | 194,176 | 0,131 | 0,9992 | [-0,193;0,214] | 0,041 | [-0,577;0,659] |
| | PI | Initial Baseline - Play | -0,071 | 0,089 | 193,048 | -0,802 | 0,8533 | [-0,301;0,159] | -0,284 | [-0,982;0,414] |
| | | Initial Baseline - Reading | -0,293 | 0,089 | 193,048 | -3,305 | 0,0062 | [-0,523;-0,063] | -1,169 | [-1,874;-0,464] |
| | | Initial Baseline - Final Baseline | -0,113 | 0,092 | 196,225 | -1,225 | 0,6119 | [-0,353;0,126] | -0,451 | [-1,179;0,277] |
| | | Play - Reading | -0,222 | 0,089 | 193,048 | -2,503 | 0,0626 | [-0,452;0,008] | -0,885 | [-1,587;-0,183] |
| | | Play – Final Baseline | -0,042 | 0,092 | 196,225 | -0,455 | 0,9686 | [-0,282;0,197] | -0,168 | [-0,895;0,559] |
| | | Reading - Final Baseline | 0,18 | 0,092 | 196,225 | 1,947 | 0,2121 | [-0,060;0,420] | 0,717 | [-0,012;1,447] |
| | T | Initial Baseline - Play | 0,021 | 0,063 | 193,048 | 0,331 | 0,9875 | [-0,142;0,183] | 0,083 | [-0,411;0,576] |

| | | | | | | | | |
|---|---|---|---|---|---|---|---|---|
| Initial Baseline - Reading | 0,02 | 0,063 | 193,048 | 0,326 | 0,988 | [-0,142;0,183] | 0,081 | [-0,412;0,575] |
| Initial Baseline - Final Baseline | 0,006 | 0,064 | 194,534 | 0,096 | 0,9997 | [-0,160;0,172] | 0,025 | [-0,479;0,528] |
| Play - Reading | 0 | 0,063 | 193,048 | -0,005 | 1 | [-0,163;0,162] | -0,001 | [-0,495;0,492] |
| Play - Final Baseline | -0,015 | 0,064 | 194,534 | -0,228 | 0,9958 | [-0,180;0,151] | -0,058 | [-0,562;0,445] |
| Reading - Final Baseline | -0,014 | 0,064 | 194,534 | -0,223 | 0,9961 | [-0,180;0,152] | -0,057 | [-0,560;0,447] |

**Table S10. Infant Behavioural analysis at test, comparison after the intervention (T1)**. Results from linear mixed-effects models fitted to parental report scores. The first column lists model terms. *Group* denotes the three infant groups (Term, Preterm Intervention, Preterm Control). *propPI* and *propPC* indicate propensity scores, modeled as described in Materials and Methods.

| ESCS | | | | |
|---|---|---|---|---|
| **IJA** | | | | |
| | *df* | *F-value* | *p-value* | η²partial |
| *Group* | 2 | 3,498 | 0,0387 | 0,129 |
| *propPI* | 1 | 0,481 | 0,4916 | 0,001 |
| *propPC* | 1 | 1,158 | 0,2877 | 0,025 |
| *Residuals* | 45 | | | |
| **RJA** | | | | |
| | *df* | *F-value* | *p-value* | η²partial |
| *Group* | 2 | 1,415 | 0,2539 | 0,047 |
| *propPI* | 1 | 8,064 | 0,0068 | 0,064 |
| *propPC* | 1 | 5,070 | 0,0294 | 0,103 |
| *Residuals* | 44 | | | |
| **IBR** | | | | |
| | *df* | *F-value* | *p-value* | η²partial |
| *Group* | 2 | 0,706 | 0,4989 | 0,036 |
| *propPI* | 1 | 1,182 | 0,2826 | 0,017 |
| *propPC* | 1 | 0,446 | 0,5075 | 0,010 |
| *Residuals* | 46 | | | |
| **RBR(%)** | | | | |
| | *df* | *F-value* | *p-value* | η²partial |
| *Group* | 2 | 0,002 | 0,9980 | 0,005 |
| *propPI* | 1 | 0,566 | 0,4558 | 4,28E-05 |
| *propPC* | 1 | 1,039 | 0,3135 | 0,022 |
| *Residuals* | 46 | | | |
| **RSI** | | | | |
| | *df* | *F-value* | *p-value* | η²partial |
| *Group* | 2 | 0,229 | 0,7963 | 0,013 |
| *propPI* | 1 | 0,596 | 0,4440 | 0,004 |
| *propPC* | 1 | 0,432 | 0,5141 | 0,009 |
| *Residuals* | 46 | | | |

**Table S11. Infant formal development at test, comparison after the intervention (T1).** Results from linear mixed-effects models fitted developmental scores. The first column lists model terms. Group denotes the three infant groups (Term, Preterm Intervention, Preterm Control). propPI and propPC indicate propensity scores, modeled as described in Materials and Methods. Assessment of infants development has been performed using the Language and the Cognitive scales Bayley Scales of Infant and Toddler Development (Bayley-III).

| Bayley | | | | |
|---|---|---|---|---|
| **Cognitive** | | | | |
| | *df* | *F-value* | *p-value* | $\eta^2$partial |
| *Group* | 2 | 0,344 | 0,7105 | 0,104 |
| *propPI* | 1 | 2,51E-04 | 0,9874 | 0,027 |
| *propPC* | 1 | 0,863 | 0,3576 | 0,018 |
| *Residuals* | 48 | | | |
| **Language** | | | | |
| | *df* | *F-value* | *p-value* | $\eta^2$partial |
| *Group* | 2 | 1,171 | 0,3186 | 0,046 |
| *propPI* | 1 | 0,136 | 0,7142 | 0,007 |
| *propPC* | 1 | 2,22E-04 | 0,9882 | 4,54E-06 |
| *Residuals* | 49 | | | |

**Table S12. Post-hoc on significant outcome - Infant Behavioural analysis at test, comparison after the intervention (T1)**. Post-hoc analysis on behavioural scores. The first column lists the group compared. (T = Term, PI = Preterm Intervention, PC = Preterm Control).

| ESCS | | | | | | | | |
|---|---|---|---|---|---|---|---|---|
| IJA | | | | | | | | |
| contrast | Mean Diff | SE | df | t-ratio | p-value | 95% CI | Cohen's d | 95% CI (d) |
| PI - PC | 0,307 | 0,122 | 45,000 | 2,522 | 0,0397 | [0,012,0,601] | 0,921 | [0,160,1,683] |
| PI - T | 0,251 | 0,143 | 45,000 | 1,761 | 0,1944 | [-0,095,0,597] | 0,756 | [-0,123,1,634] |
| PC - T | -0,055 | 0,145 | 45,000 | -0,382 | 0,9230 | [-0,406,0,295] | -0,166 | [-1,042,0,710] |

**Table S13. Parental report at test, comparison after the intervention (T1)**. Results from linear mixed-effects models fitted to parental report scores. The first column lists model terms. *Group* denotes the three infant groups (Term, Preterm Intervention, Preterm Control). *propPI* and *propPC* indicate propensity scores, modeled as described in Materials and Methods.

| **PSI** | | | | |
|---|---|---|---|---|
| **Total stress** | | | | |
| | *df* | *F-value* | *p-value* | $\eta^2$partial |
| *Group* | 2 | 0,492 | 0,6136 | 0,008 |
| *propPI* | 1 | 6,815 | 0,0110 | 0,017 |
| *propPC* | 1 | 6,024 | 0,0165 | 0,077 |
| *Residuals* | 72 | | | |
| **Parental distress** | | | | |
| | *df* | *F-value* | *p-value* | $\eta^2$partial |
| *Group* | 2 | 0,763 | 0,4702 | 0,010 |
| *propPI* | 1 | 6,776 | 0,0112 | 0,020 |
| *propPC* | 1 | 5,564 | 0,0211 | 0,072 |
| *Residuals* | 72 | | | |
| **Parent-child disfunction** | | | | |
| | *df* | *F-value* | *p-value* | $\eta^2$partial |
| *Group* | 2 | 0,339 | 0,7133 | 0,005 |
| *propPI* | 1 | 4,222 | 0,0435 | 0,002 |
| *propPC* | 1 | 5,277 | 0,0245 | 0,068 |
| *Residuals* | 72 | | | |
| **Difficult child** | | | | |
| | *df* | *F-value* | *p-value* | $\eta^2$partial |
| *Group* | 2 | 0,337 | 0,7154 | 0,013 |
| *propPI* | 1 | 3,909 | 0,0519 | 0,014 |
| *propPC* | 1 | 2,968 | 0,0892 | 0,040 |
| *Residuals* | 72 | | | |
| **RSQ** | | | | |
| **Total** | | | | |
| | *df* | *F-value* | *p-value* | $\eta^2$partial |
| *Group* | 2 | 2,893 | 0,0620 | 0,092 |
| *propPI* | 1 | 6,418 | 0,0135 | 0,001 |
| *propPC* | 1 | 9,287 | 0,0032 | 0,116 |
| *Residuals* | 71 | | | |
| **F1** | | | | |
| | *df* | *F-value* | *p-value* | $\eta^2$partial |
| *Group* | 2 | 0,326 | 0,7229 | 0,024 |
| *propPI* | 1 | 1,603 | 0,2096 | 0,008 |
| *propPC* | 1 | 1,061 | 0,3066 | 0,015 |
| *Residuals* | 71 | | | |

| F2 | | | | |
|---|---|---|---|---|
| | *df* | *F-value* | *p-value* | $\eta^2$partial |
| *Group* | 2 | 0,229 | 0,7963 | 0,012 |
| *propPI* | 1 | 11,156 | 0,0013 | 0,084 |
| *propPC* | 1 | 5,072 | 0,0274 | 0,067 |
| *Residuals* | 71 | | | |

| F3 | | | | |
|---|---|---|---|---|
| | *df* | *F-value* | *p-value* | $\eta^2$partial |
| *Group* | 2 | 6,693 | 0,0022 | 0,099 |
| *propPI* | 1 | 0,017 | 0,8969 | 0,086 |
| *propPC* | 1 | 4,718 | 0,0332 | 0,062 |
| *Residuals* | 71 | | | |

| EPDS | | | | |
|---|---|---|---|---|
| | *df* | *F-value* | *p-value* | $\eta^2$partial |
| *Group* | 2 | 1,308 | 0,2774 | 0,023 |
| *propPI* | 1 | 10,303 | 0,0021 | 0,057 |
| *propPC* | 1 | 6,361 | 0,0141 | 0,088 |
| *Residuals* | 66 | | | |

**Table S14. Post-hoc on significant outcome - Parental report at test, comparison after the intervention (T1)**. Post-hoc analysis on parental report scores. The first column lists the group compared (T = Term, PI = Preterm Intervention, PC = Preterm Control).

| RSQ | | | | | | | | |
|---|---|---|---|---|---|---|---|---|
| **Total** | | | | | | | | |
| ***contrast*** | ***Mean Diff*** | ***SE*** | ***df*** | ***t-ratio*** | ***p-value*** | ***95% CI*** | ***Cohen's d*** | ***95% CI (d)*** |
| PI - PC | 0,002 | 0,118 | 71 | 0,02 | 0,9998 | [-0,280;0,284] | 0,006 | [-0,627;0,638] |
| PI - T | 0,274 | 0,132 | 71 | 2,07 | 0,1028 | [-0,042;0,591] | 0,738 | [ 0,018;1,459] |
| PC - T | 0,272 | 0,124 | 71 | 2,20 | 0,0789 | [-0,025;0,569] | 0,733 | [ 0,056;1,410] |
| **F3** | | | | | | | | |
| ***contrast*** | ***Mean Diff*** | ***SE*** | ***df*** | ***t-ratio*** | ***p-value*** | ***95% CI*** | ***Cohen's d*** | ***95% CI (d)*** |
| PI - PC | 0,467 | 0,178 | 71 | 2,63 | 0,0279 | [ 0,042;0,892] | 0,834 | [ 0,187;1,482] |
| PI - T | 0,705 | 0,199 | 71 | 3,53 | 0,0021 | [ 0,227;1,182] | 1,258 | [ 0,518;1,999] |
| PC - T | 0,238 | 0,187 | 71 | 1,27 | 0,4161 | [-0,210;0,685] | 0,424 | [-0,245;1,094] |
